# Evaluation of Synergy Extrapolation for Predicting Unmeasured Muscle Excitations from Measured Muscle Synergies

**DOI:** 10.1101/2020.08.05.238840

**Authors:** Di Ao, Mohammad S. Shourijeh, Carolynn Patten, Benjamin J. Fregly

## Abstract

Electromyography (EMG)-driven musculoskeletal modeling relies on high-quality measurements of muscle electrical activity to estimate muscle forces. However, a critical challenge for practical deployment of this approach is missing EMG data from muscles that contribute substantially to joint moments. This situation may arise due to either the inability to measure deep muscles with surface electrodes or the lack of a sufficient number of EMG electrodes. Muscle synergy analysis is a dimensionality-reduction approach to decompose a large number of muscle excitations into a small number of time-varying synergy excitations along with time-invariant synergy weights that define the contribution of each corresponding synergy excitation to a specific muscle excitation. This study evaluates how accurately missing muscle excitations can be predicted using synergy excitations extracted from muscles with available EMGs (henceforth called “synergy extrapolation”). The results were reported on a gait dataset collected from a stroke survivor walking on an instrumented treadmill at self-selected and fastest-comfortable speeds. The evaluation process started with full calibration of a lower-body EMG-driven model using 16-channel EMGs (including surface and indwelling) in each leg. One indwelling EMG (either iliopsoas or adductor longus) was then treated as unmeasured at a time. The synergy weights associated with the unmeasured muscle were predicted through solving a nonlinear optimization problem where the errors between inverse dynamics and EMG-driven joint moments were minimized. We also quantitatively evaluated how synergy analysis algorithms (principal component analysis (PCA) and non-negative matrix factorization (NMF)), EMG normalization methods, and number of synergies affect the accuracy of the predicted unmeasured muscle excitation. Synergy extrapolation performance was most influenced by the choice of synergy analysis algorithm and number of synergies. PCA with 5 or 6 synergies consistently predicted unmeasured muscle excitations most accurately and with greatest robustness to choice of EMG normalization method. Furthermore, the associated joint moment matching accuracy was comparable to that produced by the full EMG-driven calibration. The synergy extrapolation method described in this study may facilitate the assessment of human neuromuscular control and biomechanics in response to surgical or rehabilitation treatment when important EMG signals are missing.

## 1 Introduction

Prediction of muscle forces can provide valuable insight not only for understanding the control strategies employed by the central nervous system (CNS) (Contessa and Luca, 2013; Del Vecchio et al., 2018), but also for developing effective treatments for neuromusculoskeletal disorders (Shao et al., 2009; Fregly et al., 2012b, 2012a; Allen et al., 2013; Pitto et al., 2019; Sauder et al., 2019) Since direct measurement of muscle force is invasive using current technology, computational techniques has been developed to generate muscle force estimates (Anderson and Pandy, 2001; Lloyd and Besier, 2003; Thelen et al., 2003; Buchanan et al., 2005; Shao et al., 2009) However, since the human musculoskeletal system possesses more muscles than degrees-of-freedom (DOFs) in the skeleton (i.e., the muscle redundancy problem), no unique solution for muscle exists unless muscle activity patterns are defined by measured EMG signals (Lloyd and Besier, 2003; Manal and Buchanan, 2003; Buchanan et al., 2005; Shao et al., 2009; Kumar et al., 2012; Sartori et al., 2012; Meyer et al., 2017), or assumptions are made about how muscles contribute to the joint moments, such as minimization of energetic cost (Anderson and Pandy, 2001; Ackermann and van den Bogert, 2010; S. Shourijeh and McPhee, 2014). EMG-driven musculoskeletal modeling is a computational approach for predicting muscle forces that can bypass the muscle redundancy problem while simultaneously allowing for calibration of unmeasurable musculotendon properties (e.g., optimal muscle fiber length) (Lloyd and Besier, 2003; Amarantini and Martin, 2004; Shao et al., 2009; Sartori et al., 2012; Meyer et al., 2017). In EMG-driven models, processed EMG and muscle-tendon kinematic data are input to a muscle force generation model (typically a Hill-type model) to predict muscle forces and corresponding net muscle moments. Nonlinear optimization is then used to calibrate musculotendon model parameters such that predicted net muscle moments match inverse dynamic joint moments as closely as possible.

In EMG-driven models, the quality of the measured EMG signals affects the reliability of the estimated muscle forces. Surface EMG recording, which is non-invasive and easily applicable, has been the most popular method for measuring muscle electrical activity for biomechanical studies. There are, nevertheless, intrinsic potential challenges with surface EMG data that may limit the accuracy of estimated muscle forces, such as noisy signals from crosstalk between adjacent muscles, movement artifacts, and challenges in attaining the true maximum muscle excitation for EMG normalization (Farina et al., 2002; Racinais et al., 2013; Sartori et al., 2014). Beyond these issues, the inability to acquire EMG data from deep muscles that contribute substantially to joint moments is a practical challenge (Sartori et al., 2014; Zonnino and Sergi, 2019). For instance, it is practically impossible to collect EMG data from deep hip muscles (e.g. iliacus and psoas) using surface electrodes. However, when EMG data from important deep muscles are missing in an EMG-driven model, force estimates for other muscles that have similar roles may be significantly overestimated (Zonnino and Sergi, 2019). Compared to surface electrodes, indwelling electrodes are able to measure the electrical activity of deep muscles with lower levels of crosstalk (Péter et al., 2019). However, the use of indwelling electrodes requires special skills and longer set-up time and may cause discomfort and pain for the subject. Furthermore, in some cases, such as patients who have a cancerous tumor near an important deep muscle, use an indwelling electrode for EMG data collection may be contraindicated for safety reasons. Regardless of the EMG measurement technique, EMG-driven models require EMG data collection from a large number of muscles, which may not be possible due to hardware limitation. Therefore, a computational method is needed that can provide reliably estimate muscle excitations that cannot be measured experimentally for technical or safety reasons.

Previous studies have explored computational methods for predicting unmeasured muscle excitations within the context of EMG-driven modeling. Static optimization (SO) (Crowninshield and Brand, 1981; Anderson and Pandy, 2001; Damsgaard et al., 2006; Heintz and Gutierrez-Farewik, 2007) has also been embedded into calibration of EMG-driven models, in which while the musculoskeletal model is being calibrated, missing muscle excitations can be estimated. The objective function for such technique includes minimizing both activation levels of unmeasured muscles and joint moment tracking errors (Sartori et al., 2014; Zonnino and Sergi, 2019). Zonnino et al. presented an EMG-driven forward dynamics estimator that used an SO-based neural model to determine unmeasured muscle activations, which reduced the estimation error of muscle forces in comparison to the performance of a conventional estimator that neglected the contribution of unmeasured muscles (Zonnino and Sergi, 2019). Similarly, Satori et al. developed a hybrid EMG-informed model in which experimental EMG signals were minimally adjusted while missing EMG signals (e.g., from iliacus and psoas) were predicted via SO. However, none of these studies have shown evidence that the reproduced unmeasured muscle activations are reliable and in reasonable agreement with experimental measurements. Furthermore, because time histories are not taken into account in SO, the resulting muscle activations might contain abrupt unrealistic changes due to the optimization problem being solved at an instant in time.

Another approach for estimating unmeasured muscle excitations is to use muscle synergy concepts. A muscle synergy is composed of a time-varying synergy excitation and a corresponding time-invariant synergy vector containing weights that define how each synergy excitation contributes to the excitation of each muscle (Tresch et al., 1999; Ting and Chvatal, 2010; Banks et al., 2017; Shourijeh and Fregly, 2020) Muscle synergies have been broadly used in descriptive research to analyze experimental muscle excitations during a large number of movement tasks (Ivanenko et al., 2005; Torres-Oviedo and Ting, 2007; Bowden et al., 2010; Walter et al., 2014; Kristiansen et al., 2015; Meyer et al., 2016a; Ruiz Garate et al., 2017; Sauder et al., 2019), but few studies have performed predictive analyses using muscle synergy information (Ajiboye and Weir, 2009; Bianco et al., 2018). Ajiboye and Weir demonstrated that subject-specific synergies extracted from muscle activities recorded for a subset of postures can be used to predict EMG patterns for the remaining postures. Bianco et al. investigated the theoretical feasibility of using synergy excitations extracted from a group of 8 ‘included’ muscle excitations treated as measured as building blocks to construct muscle excitations for a group of 8 ‘excluded’ muscle excitations treated as unmeasured. However, the muscle synergy weights associated with ‘remaining’ postures in (Ajiboye and Weir, 2009) or ‘excluded’ muscles in (Bianco et al., 2018) were not predicted without knowledge of the muscle excitations treated as unmeasured but rather fitted with knowledge of those excitations using least square algorithms. Several other studies have imposed synergy structures on muscle excitations or activations through optimization when estimating knee contact force (Walter et al., 2014), joint stiffness (Shourijeh and Fregly, 2020), or motion (Clark et al., 2009; Allen and Neptune, 2012; Meyer et al., 2016b; Mehrabi et al., 2019; Falisse et al., 2020). However, to the best of the authors’ knowledge, the reliability with which unmeasured muscle excitations can be estimated using EMG-driven models with unknown synergy weights has not been studied previously. Additionally, synergy extrapolation has never been evaluated from a predictive perspective, and it is therefore unknown how methodological choices for how synergy analysis is performed affect the outcome of synergy extrapolation.

A synergy-based muscle excitation estimation method involving EMG-driven modeling, termed “synergy extrapolation,” was developed to predict muscle excitations that cannot be measured experimentally. This study evaluated how reliably synergy extrapolation is able to predict missing indwelling EMG signals using a well-calibrated EMG-driven lower-body musculoskeletal model. Since the outcome of muscle synergy analysis is affected by methodological choices, such as EMG processing (e.g. magnitude normalization), physiological assumptions (e.g. number of synergies), and numerical analysis approaches (e.g. matrix decomposition algorithm (Tresch et al., 2006; Hug et al., 2012; Steele et al., 2013; Oliveira et al., 2014; Banks et al., 2017; Shuman et al., 2017; Ebied et al., 2018; Gallina et al., 2018)), we also evaluated how each of these choices affects synergy extrapolation results. The evaluated methodological decisions for MSA process included matrix factorization algorithm (PCA and NMF), EMG normalization approach, and number of synergies. To validate this approach, we analyzed a gait data set collected from a high-functioning stroke subject performing treadmill walking at self-selected and fastest-comfortable speeds. Muscle excitations that were measured using indwelling EMG electrodes, i.e., from iliopsoas and adductor longus, were assumed unknown and were predicted. By quantitatively evaluating the differences in synergy extrapolation performance produced by different methodological decisions, this work provides evidence-based suggestions for which methods are likely to produce the most accurate predictions of missing muscle excitations.

## 2 Materials and Methods

### 2.1 Experimental data

A previously published gait dataset collected from one high-functioning stroke survivor (age 79 years, LE Fugl-Meyer Motor Assessment 32/34 pts, right-sided hemiparesis, height 1.7 m, mass 80.5 kg) was used to evaluate the synergy extrapolation process (Meyer et al., 2017). Motion capture (100 Hz, Vicon Corp., Oxford, UK), ground reaction force (1000 Hz, Bertec Corp., Columbus, OH), and surface EMG (1000 Hz, Motion Lab Systems, Baton Rouge, LA) data were recorded simultaneously while the subject walked on a split-belt instrumented treadmill (Bertec Corp., Columbus, OH) at two speeds: 0.5 m/s (self-selected speed) and 0.8 m/s (fastest comfortable speed). Ten cycles of gait data for each walking speed were analyzed. Motion capture and ground reaction force (GRF) data were low-pass filtered using a fourth-order zero-phase lag Butterworth filter with a cut-off frequency at 7/*tf* Hz, where *tf* is the period of the gait cycle being processed (McLean et al., 2005; Meyer et al., 2017). Sixteen channels of EMG data were collected from each leg using surface and a small number of indwelling electrodes (i.e. fine-wire), which made available EMG data from important deep muscle groups (e.g. iliopsoas) (supplementary table 1). After being high-pass filtered at 40 Hz, demeaned, full-wave rectified, and low-pass filtered at 3.5/*tf* Hz, each EMG signal was normalized to the maximum value over all trials. After processing, each trial of EMG data was time-normalized to 101 time frames in each gait cycle (heel strike to heel strike). See (Meyer et al., 2017) for more details.

### 2.2 Musculoskeletal model

A generic full-body OpenSim musculoskeletal model (Arnold et al., 2010) was adopted for analyses in OpenSim v3.3 (Delp et al., 2007; Seth et al., 2018). The model controlled 5 degrees of freedom (DOFs), including two-DOF hip joints (flexion/extension and adduction/abduction), one DOF knee joints (flexion/extension), and two-DOF ankle joints (plantarflexion/dorsiflexion and inversion/eversion). Of the 45 muscles in each leg accounted for in the original model, 35 muscles per leg were retained. Compartments of large muscles shared common EMG signals (Meyer et al., 2017). To personalize the model, we performed five steps sequentially on the modified generic OpenSim model using a combination of OpenSim and custom Matlab analyses: 1) model scaling to match the subject’s anthropometry; 2) kinematic calibration to determine personalized lower body joint positions and orientations (Reinbolt et al., 2005); 3) inverse kinematics (IK) to calculate joint angle time histories from the surface marker data using the calibrated kinematic model; 4) surrogate musculoskeletal geometry creation to fit muscle-tendon lengths and moment arms as polynomial functions of lower body joint angles and velocities (Menegaldo et al., 2004; Meyer et al., 2017); and 5) inverse dynamics (ID) to obtain experimental joint moments using experimental GRF data and IK-derived time histories of joint kinematics as inputs.

### 2.3 Synergy extrapolation methodology

Our proposed synergy extrapolation method consists of two functional blocks (Fig. 1). First, the synergy extrapolation block seeks to reconstruct unmeasured muscle excitations with measured synergy excitations and a set of unmeasured synergy vector weights that is found through an optimization process. Muscle synergy analysis is performed trial by trial for all measured muscle excitations ***e***_***m***_ to extract a low-dimensional set of time-varying measured synergy excitations ***W***_***m***_ and corresponding sets of time-invariant measured weighting vectors ***H***_***m***_, where each weighting vector represents the contribution of the corresponding synergy excitation to all measured muscle excitations. Next, unmeasured muscle excitations ***e***_***x***_ are constructed using the measured synergy excitations ***W***_***m***_ along with a set of trial-specific synergy vector weights ***H***_***x***_ associated with the unmeasured muscles and predicted by the optimization. In the present study, several methodological assumptions and decisions were made prior to MSA, including number of synergies, synergy analysis approach, and EMG normalization method. The key differences in the implementation of these different methodologies for synergy extrapolation are described below. Second, the EMG-driven musculoskeletal model block predicts the net joint moments generated by muscles using a combination of experimentally measured muscle excitations (***e***_***m***_) and computationally predicted muscle excitations (***e***_***x***_) constructed from measured synergy excitations ***W***_***m***_ and predicted synergy vector weights ***H***_***x***_. In this block, a previously developed and calibrated EMG-driven model, where muscles were treated as Hill-type models (Hill, 1938; Zajac, 1989) with a rigid tendon, was employed to predict muscle forces and net joint moments in the lower extremities (Meyer et al., 2017). The joint moment produced by a muscle spanning a particular joint can be represented as:

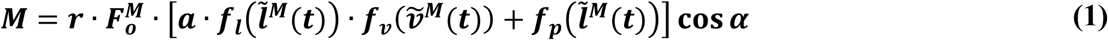

where ***M*** is the moment generated by the muscle about the joint, ***r*** is the moment arm of the muscle about the same joint, 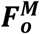 is the maximum isometric force of the muscle, ***a*** is the muscle activation, 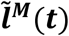 and 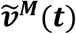 are the time-varying normalized muscle fiber length and velocity, respectively, and *a* is the pennation angle of the muscle. 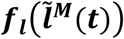 and 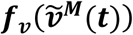 define the normalized muscle active force-length and active force-velocity relationships, while 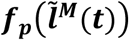 defines the normalized muscle passive force-length relationship (Zajac, 1989)(Meyer et al., 2017). Muscle activation (***a***) was calculated from muscle excitation (***e***) using a model of muscle activation dynamics (He et al., 1991; Lloyd and Besier, 2003; Meyer et al., 2017). Muscle excitations were derived by multiplying the processed EMG signals by muscle-specific scale factors between 0.05 and 1 to reflect unknown maximum excitation levels.

**Fig. 1.**
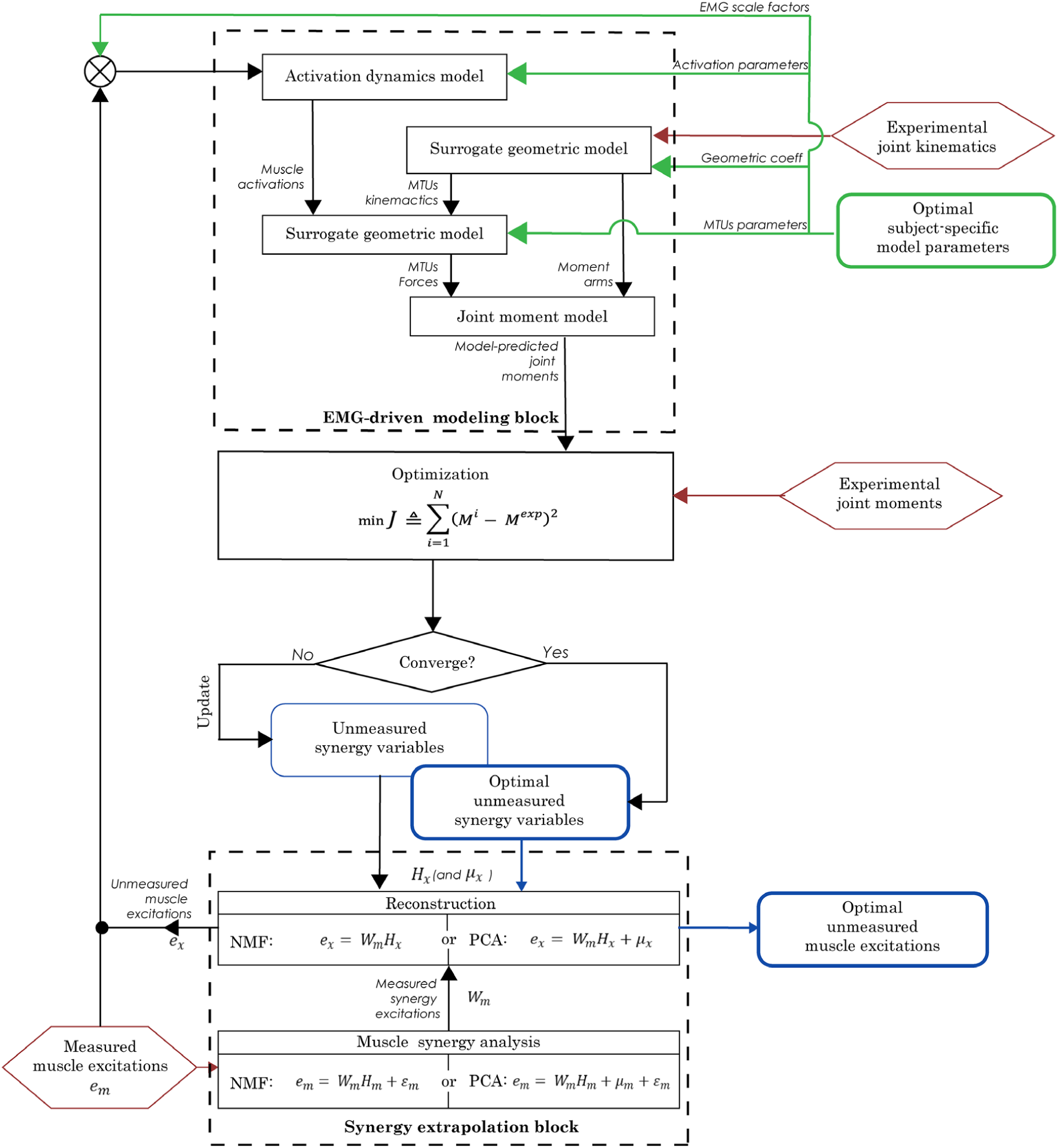
Flowchart of EMG-driven model-based synergy extrapolation framework. The framework comprises of two main blocks: synergy extrapolation block and EMG-driven musculoskeletal model block. In the synergy extrapolation block, measured muscle excitations are decomposed to determine a set of measured synergy excitations, which are used as building blocks to construct unmeasured muscle excitations. In the EMG-driven model block, the joint moments are predicted by receiving both the measured muscle excitations from experimental measurements along with the unmeasured muscle excitations as the output of the synergy extrapolation block. The unmeasured synergy variables were calibrated through the optimization process with an objective function of minimizing the difference between experimental and model-predicted joint moments. For NMF-based synergy extrapolation, the unmeasured synergy variables include the unmeasured synergy weight vectors, while for PCA-based synergy extrapolation, the unmeasured synergy variables also include an additional set of design variables that represent mean values of unmeasured muscle excitations.

In this study, unmeasured muscle excitations were predicted by following a two-step process. The first step calibrated an EMG-driven model of each leg using a full set of 16 EMG channels per leg (henceforth called ‘full EMG-driven’), where every muscle included in the EMG-driven model was associated to an experimentally measured EMG signal. During this step, a sequence of optimizations was performed to identify the parameter values required by the activation dynamics models, Hill-type muscle-tendon models, and surrogate musculoskeletal geometric model (described in Meyer et al., 2017) that reproduced the lower body inverse dynamic joint moments as closely as possible (Eq. 2). The model parameter values calibrated for each muscle-tendon actuator by the optimization process included: electromechanical delay, activation time constant, activation nonlinearity constant, EMG scale factor, optimal muscle fiber length, tendon slack length, and geometric coefficients defining muscle-tendon lengths, velocities, and moment arms. The second step calibrated the synergy vector weights, ***H***_***x***_ for the unmeasured muscle excitations using the EMG-driven model parameters found in the first step. During this step, only unmeasured synergy vector weights ***H***_***x***_ were identified iteratively through optimization. Thus, this two-step process estimated missing muscle excitations assuming a well-calibrated EMG-driven model was already available, including calibrated parameter values for the muscles with missing excitations. Predicting missing muscle excitations reliably using a well-calibrated EMG-driven model is an important step toward predicting missing muscle excitations while simultaneously calibrating the parameter values of the EMG-driven model. The structure of unmeasured synergy variables was slightly different between the matrix factorization algorithms, and more details are provided in the section of methodological choices on synergy extrapolation.

The primary cost function for both calibration steps was formulated as:

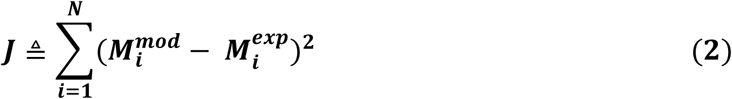

where 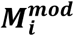 is the model-predicted moment about joint *i*, 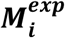 is the experimental moment about joint *i* calculated using inverse dynamics, and ***N*** is the total number of joints. Further details on the specification of initial guesses, variable bounds, overall cost function structure, additional constraints, and penalty terms for calibrating the EMG-driven model can be found in (Meyer et al., 2017). For the second step, the missing synergy vector weights, ***H***_***x***_, were unbounded and were initialized by the optimization using randomly chosen values between 0 and 1. However, the unmeasured muscle excitations predicted by synergy extrapolation were constrained to be between 0 and 1. All calibrations were performed using Matlab’s built-in “fmincon” optimization algorithm using sequential quadratic programming.

### 2.4 Methodological choices for synergy extrapolation

Muscle synergy analysis requires a number of methodological choices that can influence the results of the analysis (Banks et al., 2017). Methodological choices that have been studied include EMG processing approaches (e.g. filtering parameters, normalization methods), assumptions about neural control complexity (e.g. number of synergies, number and choice of muscles, synergy vector variability across trials), numerical analysis approaches (e.g., matrix decomposition algorithm, selection criteria for number of synergies), and post-processing of results (Tresch et al., 2006; Hug et al., 2012; Steele et al., 2013; Oliveira et al., 2014; Shourijeh et al., 2016; Banks et al., 2017; Shuman et al., 2017; Ebied et al., 2018; Gallina et al., 2018; Mehryar et al., 2020). In the present study, for each subset of measured muscles, we performed synergy extrapolation using a total of 80 methodological combinations comprised of 2 algorithms for matrix factorization, 5 methods for EMG normalization, and 8 choices for number of muscle synergies.

The two matrix factorization algorithms that render the most divergent results for MSA were used to calculate measured muscle synergies: non-negative matrix factorization (NMF) and principal component analysis (PCA) (Olree and Vaughan, 1995; Lee and Seung, 1999; d’Avella et al., 2003; Tresch et al., 2006; Ting and Chvatal, 2010; Banks et al., 2017; Bianco et al., 2018; Ebied et al., 2018). NMF is a nonlinear decomposition method, the solution of which is found iteratively to minimize the error between the reconstructed and the original dataset subject to constraints on non-negativity. Typically, NMF derives components that are similar across multiple decompositions but not numerically unique due to the non-convexity of its search space (Shourijeh et al., 2016). In contrast, PCA is a linear analytical operation that seeks to identify the internal structure of the data in a way that best explains its variance. PCA creates a unique solution of orthogonal principal components composed of both positive and negative values (Torres-Oviedo and Ting, 2007; Ting and Chvatal, 2010). During synergy extrapolation with a given number of synergies, muscle excitations in the measured muscle subset ***e***_***m***_ were represented as:

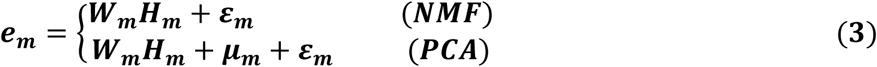

where ***e***_***m***_ denotes an ***n*** time points × 16 measured EMG signals matrix that contains measured muscle excitations in columns, ***W***_***m***_ denotes an ***n*** time points × ***p*** synergies matrix that contains measured synergy excitations in columns, and ***H***_***m***_ denotes a *p* synergies *x* 16 measured EMG signals matrix that contains measured synergy vector weights. In preparation for NMF or PCA, measured EMG signals from each gait cycle were re-sampled to 101 time frames plus 10/*tf* time frames before the start of the cycle to account for a maximum electromechanical delay of 100 ms. For NMF and PCA, ***ε***_***m***_ denotes the part of ***e***_***m***_ that cannot be explained by ***W***_***m***_***H***_***m***_, while for PCA, ***µ***_***m***_ specifies the average muscle excitations in ***e***_***m***_. Matlab functions “nnmf” (alternating least squares algorithm with 10 replicates) and “pca” were used to perform NMF and PCA, respectively.

Following MSA by either approach, unmeasured muscle excitations ***e***_***x***_ were constructed from the measured synergy excitations ***W***_***m***_ using the relationships below:

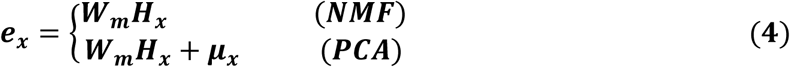

where ***H***_***x***_ is a *p*-synergy × 1 vector representing the unmeasured muscle synergy vector weights. Unlike NMF, PCA needs an additional design variable ***µ***_***x***_ that represents the average value of each unmeasured muscle excitation. Both ***H***_***x***_ and ***µ***_***x***_ were calibrated by tracking experimental joint moments within our EMG-driven modeling framework (Eq. 2).

Because EMG normalization method affects the results of MSA, this study explored five approaches for normalizing the magnitudes of processed EMG signals in preparation for MSA. EMG normalization was performed either within individual trials (Per trial) or across all trials (Over all trials). Specifically, each measured muscle EMG signal was normalized using: 1) maximum value over all trials (MaxOver), 2) maximum value per trial (MaxPer), 3) unit variance over all trials (VarOver), 4) unit variance per trial (VarPer), and 5) unit magnitude per trial (MagPer) (Banks et al., 2017). VarOver and VarPer normalizations involved dividing each EMG signal by its standard deviation over all trials and in each trial, respectively. MagPer normalization involved dividing each EMG signal by its 2-norm value for each trial (Banks et al., 2017).

Since the specified number of muscle synergies also affects the outcome of MSA substantially, we repeated the synergy extrapolation process for three through ten synergies. This range was chosen since three to six muscle synergies have been shown sufficient to account for over 90% of the variability in up to thirty muscle excitations during human movement (Ivanenko et al., 2005; Ting and Macpherson, 2005; Cappellini et al., 2006; Bianco et al., 2018). Performance of both synergy extrapolation and joint moment tracking was taken into account when determining the optimal number for synergies.

### 2.5 Evaluation of synergy extrapolation performance

To implement the first step of the synergy extrapolation procedure (full EMG-driven calibration), we randomly selected 10 gait cycles from the two walking speeds (five trials per speed) (henceforth called “calibration trials”) to calibrate the subject-specific EMG-driven musculoskeletal model. In preparation for synergy extrapolation, we assumed one hip muscle EMG signal at a time (either iliopsoas or adductor longus) recorded using indwelling electrodes to be unmeasured while the remaining 15 channels of EMG data were treated as measured. Using the calibrated full EMG-driven model, we then ran synergy extrapolation on the 10 trials used for calibration as well as on another 10 trials (5 trials per speed) randomly selected from the remaining gait cycles (henceforth called “evaluation trials”).

Several common metrics were employed to score outcomes of muscle synergy analysis, performance of synergy extrapolation, and accuracy of joint moment estimates with different combinations of methodological choices. Variance accounted for (VAF) was calculated to compare the ability of methodological combinations to construct measured muscle excitations (Shourijeh et al., 2016). Root mean square error (RMSE) and Pearson correlation coefficient *r* between mean experimental and model-predicted muscle excitations across all trials were computed to assess quantitatively matching of magnitude and shape, respectively. In addition, RMSE and *r* values were calculated for each trial, and the frequency of number of synergies with which highest *r* values or lowest RMSE values appeared was analyzed. The correlation between predicted and experimental unmeasured muscle excitations was interpreted quantitatively as weak (*r* <0.35), moderate (0.35< *r* ≤0.67), strong (0.67< *r* ≤0.9), and very strong (*r* ≥0.9) (Taylor, 1990). Mean absolute errors (MAE) between experimental and model-predicted joint moments were calculated across all gait cycles to evaluate the accuracy of the predicted joint moments during full EMG-driven calibration (step 1) and synergy extrapolation (step 2).

### 2.6 Statistical analyses

Multiple statistical analyses were used to assess whether the calculated metrics resulting from different synergy extrapolation methodological choices were statistically different for each unmeasured muscle-leg combination. First, to assess whether reconstruction performance of measured muscle excitations was statistically different between the two matrix factorization algorithms and the five EMG normalization methods, we performed a two-factor ANOVA with a Tukey-Kramer post-hoc analysis on VAF values. Second, to compare synergy extrapolation performance for different methodological choices, we conducted two three-factor (matrix factorization algorithm by EMG normalization method by number of synergies) ANOVA tests on *r* and RMSE values between predicted and experimental unmeasured muscle excitations across all calibration and evaluation trials, respectively. In addition, we also performed paired *t*-tests on *r* and RMSE values to investigate whether matrix factorization algorithm (i.e., PCA and NMF) had a significant influence on synergy extrapolation performance for the same number of synergies. Third, a three-factor (matrix factorization algorithm by EMG normalization method by number of synergies) ANOVA was carried out to compare MAE values characterizing the accuracy of joint moment tracking from different approaches. All statistical analyses were performed in Matlab, and significance levels were set at *p ≤* 0.05.

## 3 Results

### 3.1 Muscle synergy analysis

The two-way ANOVA for mean VAF values revealed main effects of matrix factorization algorithm (p < 0.01) and EMG normalization method (*p* < 0.01) on the variance explained by factorization of measured muscle excitations. For five or fewer synergies, PCA generally had significantly higher VAF values than did NMF for the same number of synergies (all *p* values < 0.05, grey shading in Table 1). Overall, extracted synergy excitations were able to predict the measured muscle excitations with > 90% VAF using 3 or more synergies in the left leg and 4 or more synergies in the right leg for PCA and 4 or more synergies in both legs for NMF (Table 1). Across all EMG normalization methods, MaxOver produced significantly higher VAF values than did MaxPer (*p* < 0.01), VarPer (*p* = 0.028), and MagPer (*p* < 0.01), while MaxPer produced the lowest VAF amongst the six EMG normalization methods (*p* < 0.01).

**Table 1.** mean VAF values across all trials that represent reconstruction quality of measured muscle excitations using 3 to 7 synergies and 6 EMG normalization methods when iliopsoas or adductor longus EMG channel is assumed to be unmeasured (The VAF values represented by PCA/NMF; grey shading represents a significant difference between PCA and NMF with matched EMG normalization method and matched number of synergies, P ≤ 0.05).

| Unmeasured muscle | EMG normalization | Left leg (non-paretic) |  |  |  |  | Right leg (paretic) |  |  |  |  |
| --- | --- | --- | --- | --- | --- | --- | --- | --- | --- | --- | --- |
|  |  | Number of synergies |  |  |  |  | Number of synergies |  |  |  |  |
|  |  | 3 | 4 | 5 | 6 | 7 | 3 | 4 | 5 | 6 | 7 |
| Iliopsoas | MaxOver | 92.7/88.8 | 97.7/95.4 | 99.1/98.0 | 99.7/98.8 | 99.9/99.3 | 89.7/82.3 | 95.2/91.0 | 98.1/95.4 | 99.2/97.5 | 99.7/98.5 |
|  | MaxPer | 90.8/86.5 | 97.2/94.7 | 98.8/97.5 | 99.5/98.5 | 99.8/99.1 | 85.6/77.7 | 93.1/86.8 | 97.2/93.1 | 98.7/96.3 | 99.5/97.6 |
|  | VarOver | 90.1/86.4 | 97.4/94.4 | 98.9/97.6 | 99.6/98.5 | 99.9/99.0 | 87.8/77.8 | 94.6/89.1 | 97.8/94.1 | 99.1/96.4 | 99.7/97.6 |
|  | VarPer | 90.9/87.3 | 97.2/94.6 | 98.8/97.5 | 99.5/98.5 | 99.8/99.1 | 85.7/77.9 | 93.3/86.6 | 97.2/92.7 | 98.7/96.2 | 99.5/97.8 |
|  | MagPer | 92.5/88.5 | 97.4/95.0 | 98.9/97.7 | 99.6/98.6 | 99.9/99.2 | 83.5/75.6 | 91.9/85.9 | 96.7/91.5 | 98.8/95.6 | 99.5/97.6 |
| Adductor longus | MaxOver | 94.0/90.1 | 97.9/95.8 | 99.0/97.8 | 99.7/98.8 | 99.9/99.3 | 88.8/81.3 | 94.3/90.5 | 97.5/94.8 | 99.0/97.2 | 99.7/98.5 |
|  | MaxPer | 91.9/87.8 | 97.2/94.7 | 98.7/97.2 | 99.5/98.4 | 99.8/99.0 | 85.1/77.2 | 91.7/86.4 | 96.5/92.6 | 98.4/95.7 | 99.5/97.5 |
|  | VarOver | 93.2/87.6 | 97.5/94.6 | 98.8/97.4 | 99.6/98.4 | 99.9/99.0 | 86.1/78.0 | 93.2/88.6 | 96.8/93.3 | 98.6/95.5 | 99.6/97.5 |
|  | VarPer | 92.4/88.3 | 97.3/94.7 | 98.7/97.2 | 99.5/98.5 | 99.8/99.0 | 84.7/76.5 | 92.1/86.1 | 96.5/92.1 | 98.5/95.8 | 99.5/97.6 |
|  | MagPer | 92.0/87.7 | 96.9/94.3 | 98.6/97.1 | 99.5/98.2 | 99.8/98.8 | 83.0/74.4 | 90.8/85.1 | 96.3/90.4 | 98.4/95.1 | 99.5/96.7 |

### 3.2 Synergy extrapolation performance

In the results for both calibration (Fig. 2) and evaluation (Fig. 3) trials, mean predicted unmeasured muscle excitations using PCA were strongly correlated to the experimental muscle excitations (mean *r* always ≥ 0.7), which was not consistently observed for the NMF results. Furthermore, RMSE values between the average predicted and actual unmeasured muscle excitations across all trials using PCA-based synergy extrapolation were generally lower than those using NMF-based synergy extrapolation. In addition, synergy extrapolation performed using either PCA or NMF predicted more accurate unmeasured muscle excitations with less trial-to-trial variability for the left leg than for the right leg (Figs. 2 and 3).

**Fig. 2.**
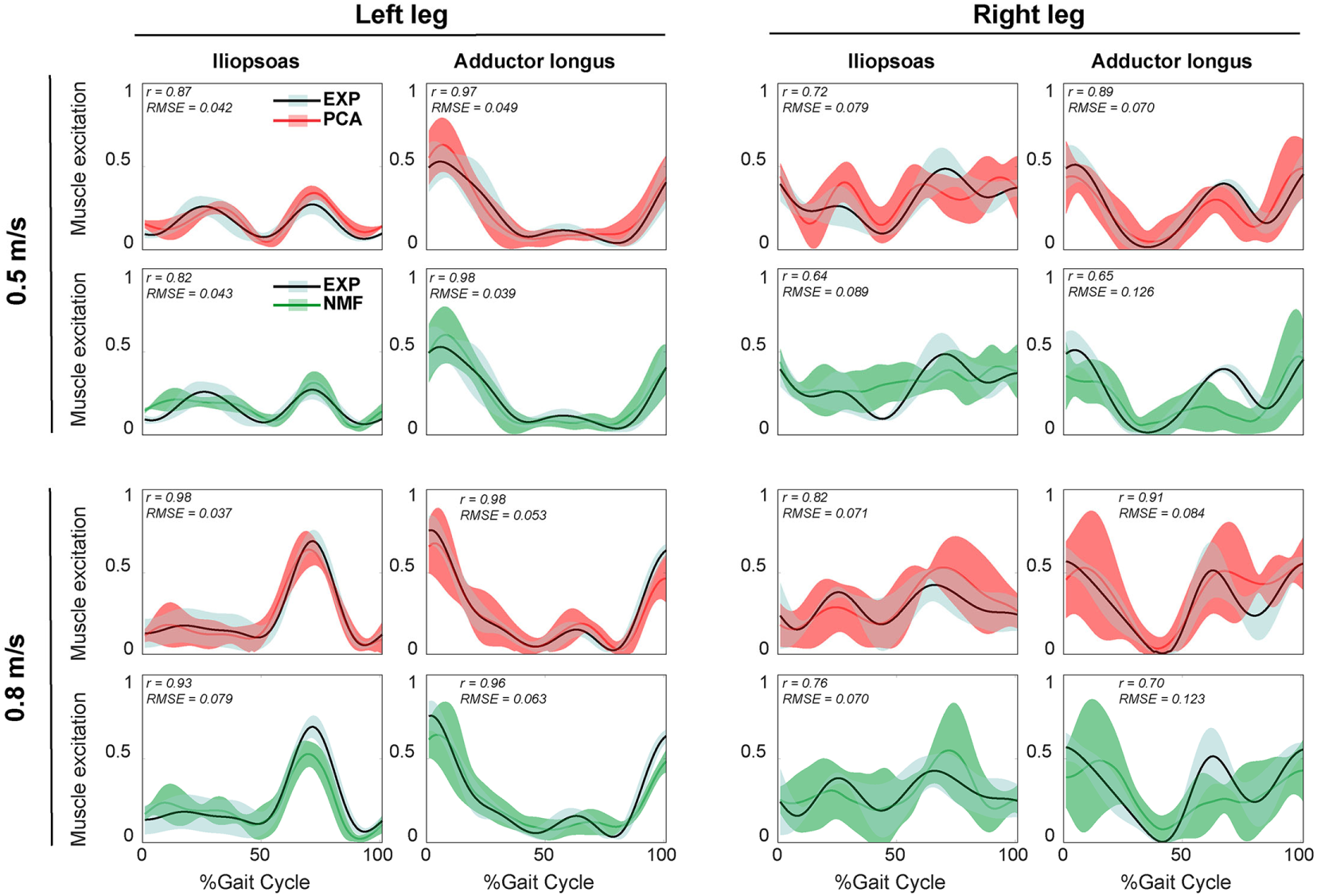
Representative results of reconstructed unmeasured muscle excitations across all calibration walking trials at the same speed using synergy extrapolation framework (black line: average experimental curve; red line: PCA-based synergy extrapolation; green line: NMF-based synergy extrapolation; shade area representing ± 1 standard deviation). The measured synergies were extracted using MaxOver EMG normalization method with 6 synergies. Dataset is reported for the whole gait cycle with 0% being heel-strike and 100% being consecutive heel-strike events for both legs (right leg: paretic, left leg: non-paretic). *r* and RMSE values are computed between average experimental and synergy-extrapolated muscle excitations.

**Fig. 3.**
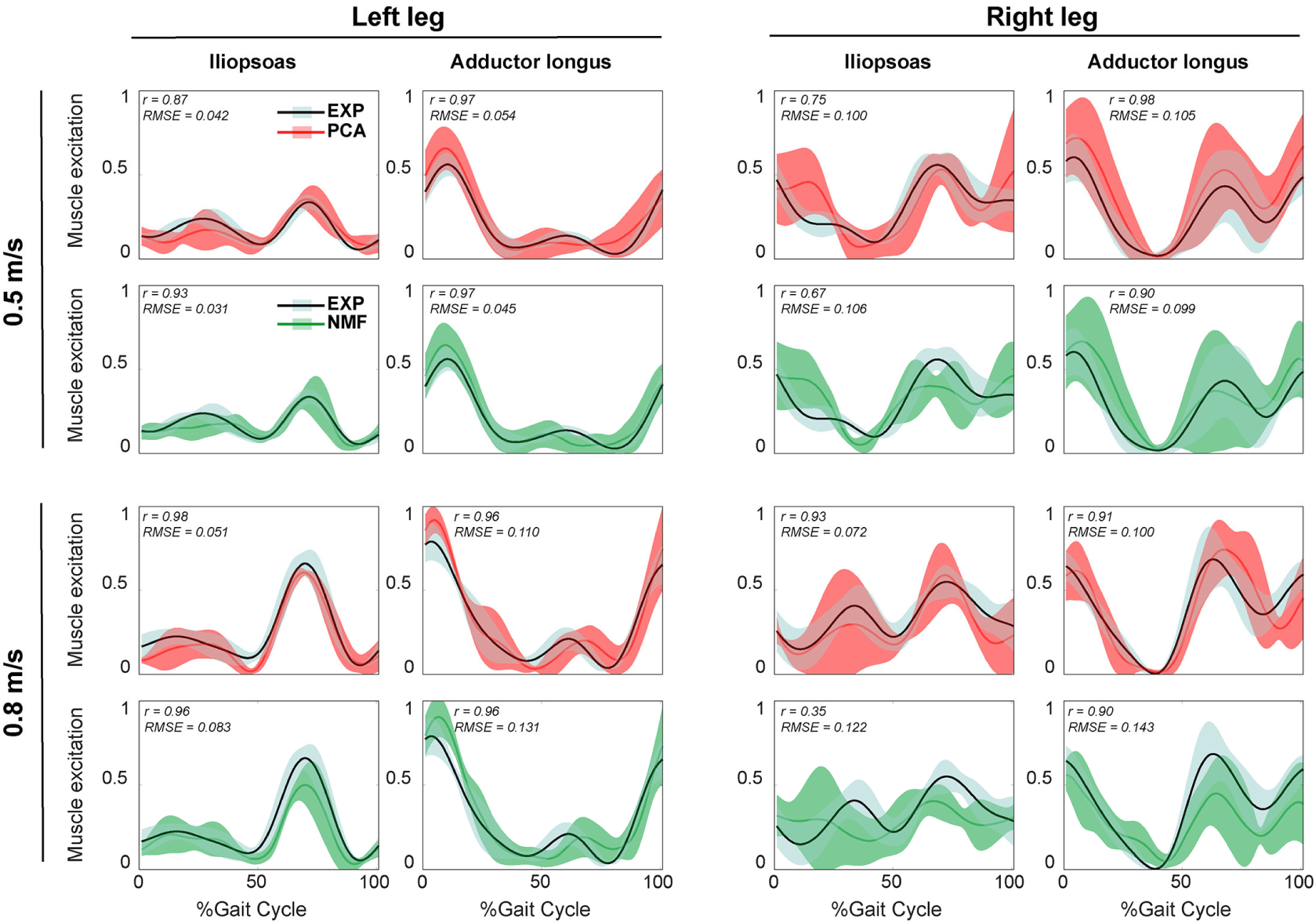
Representative results of average reconstructed unmeasured muscle excitations across all evaluation walking trials at the same speed using synergy extrapolation framework (black line: average experimental curve; red line: PCA-based synergy extrapolation; green line: NMF-based synergy extrapolation; shade area representing ± 1 standard deviation). The measured synergies, in this case, were extracted using MaxOver EMG normalization method with 6 synergies. Dataset is reported for the whole gait cycle with 0% being heel-strike and 100% being consecutive heel-strike events for both legs (right leg: paretic, left leg: non-paretic). *r* and RMSE values are derived between average experimental and synergy-extrapolated muscle excitations.

The three-factor ANOVA analyses indicated that the number of synergies (*p* < 0.01) and matrix factorization algorithm (*p* < 0.01) had a significant impact on both *r* and RMSE values for all unmeasured muscle-leg combinations, while EMG normalization method did not. For each unmeasured muscle-leg combination, as the number of synergies increased. PCA produced non-monotonic changes in *r* and RMSE values, with *r* values reaching a maximum and RMSE values reaching a minimum at 5 or 6 synergies. Unlike PCA, *r* values for NMF initially rose with an increasing number of synergies and then remained high with further increases, while RMSE values initially dropped and then leveled off (Fig. 4). Moreover, for the same number of synergies, PCA generally exhibited less variance than NMF in mean *r* and RMSE values across the six EMG normalization methods (Fig. 4).

**Fig. 4.**
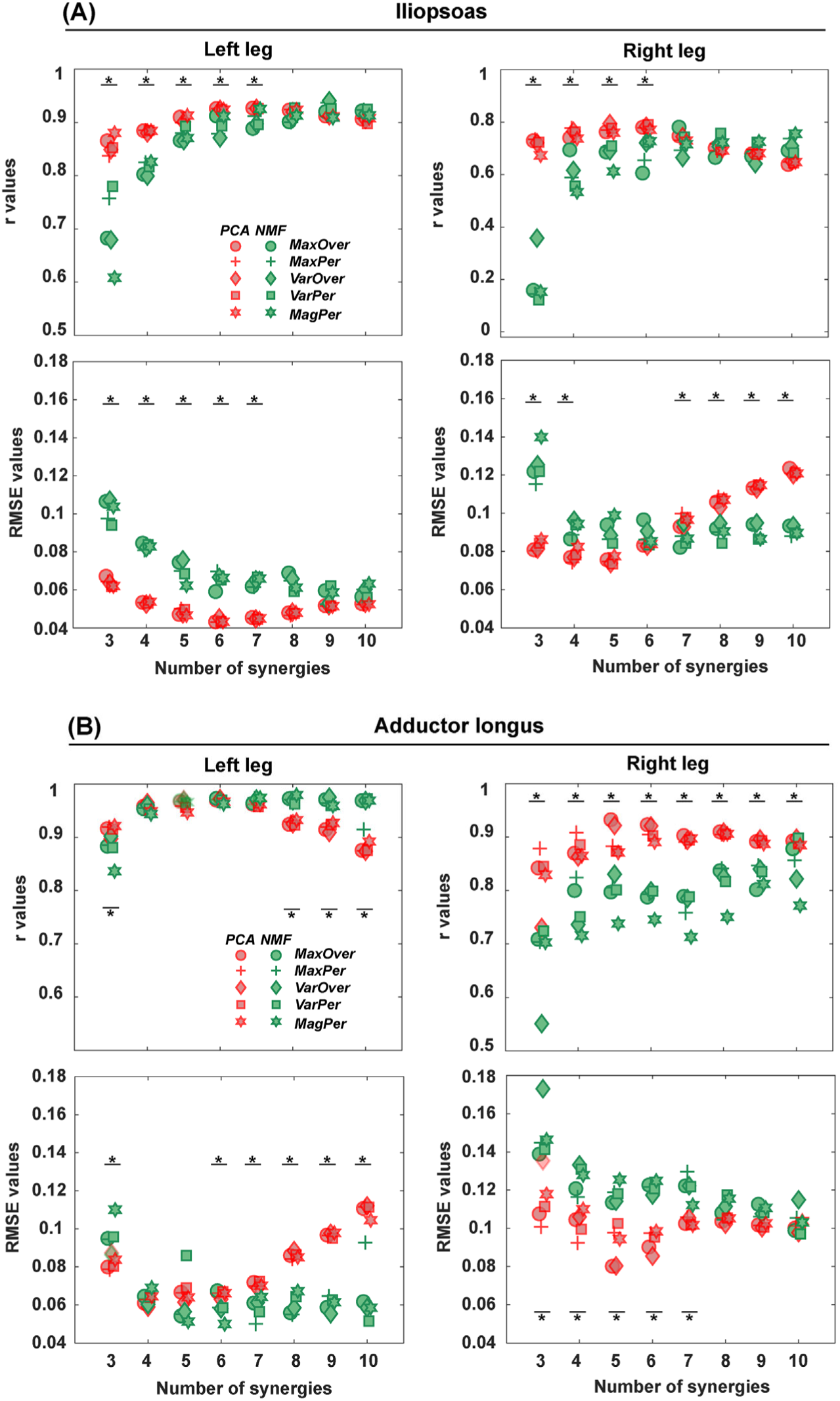
Average *r* and RMSE values for the reconstruction of iliopsoas (A) and adductor longus (B) muscle excitations across all trials (including both calibration trials and evaluation trials) for both legs (right leg: paretic, left leg: non-paretic) with 3 to 10 synergies (red markers: PCA-based synergy extrapolation; green markers: NMF-based synergy extrapolation; 5 EMG normalization methods are represented by different marker shapes; black bars represent a significant difference between PCA and NMF when matched for the number of synergies, *p* ≤ 0.05)

PCA offered significantly higher *r* values and lower RMSE values than NMF when the number of synergies varied from 3 to 6. The one exception was adductor longus in the left leg, where the only significant difference between PCA and NMF occurred for 3 synergies (*p* < 0.01). When the number of synergies increased above 6, NMF provided substantially higher r values for adductor longus in the left leg (Fig. 4 (A)), whereas PCA had markedly higher r values for adductor longus in the right leg (p < 0.05, Fig. 4 (B)). Additionally, for the number of synergies above 6, RMSE values using NMF were significantly smaller than those using PCA for iliopsoas in the right leg (Fig. 4 (A)) and for adductor longus in the left leg (Fig. 4 (B)). When similarity in both shape and magnitude are taken into account, based on average *r* and RMSE values, the best number of synergies for predicting unmeasured excitations using PCA was 6 for iliopsoas in the left leg (*r* = 0.93; RMSE = 0.043), 5 for iliopsoas in the right leg (*r* = 0.79; RMSE =0.073), 5 for adductors longus in the left leg (*r* = 0.97; RMSE =0.06), and 5 for adductors longus in the right leg (*r* = 0.93; RMSE = 0.081). Additionally, for most trials, PCA required fewer synergies (between 3 and 8) than did NMF (between 4 and 10) to achieve the best synergy extrapolation performance (see the histogram in Fig. 5). For example, for iliopsoas in the left leg, with 5 to 8 synergies, 86% of trials could achieve the highest *r* values with PCA and only 54% trials could do with NMF. Similarly, with 5 to 8 synergies, the smallest RMSE values could be obtained for 100% of trials using PCA but only 50% of trials using NMF.

**Fig. 5.**
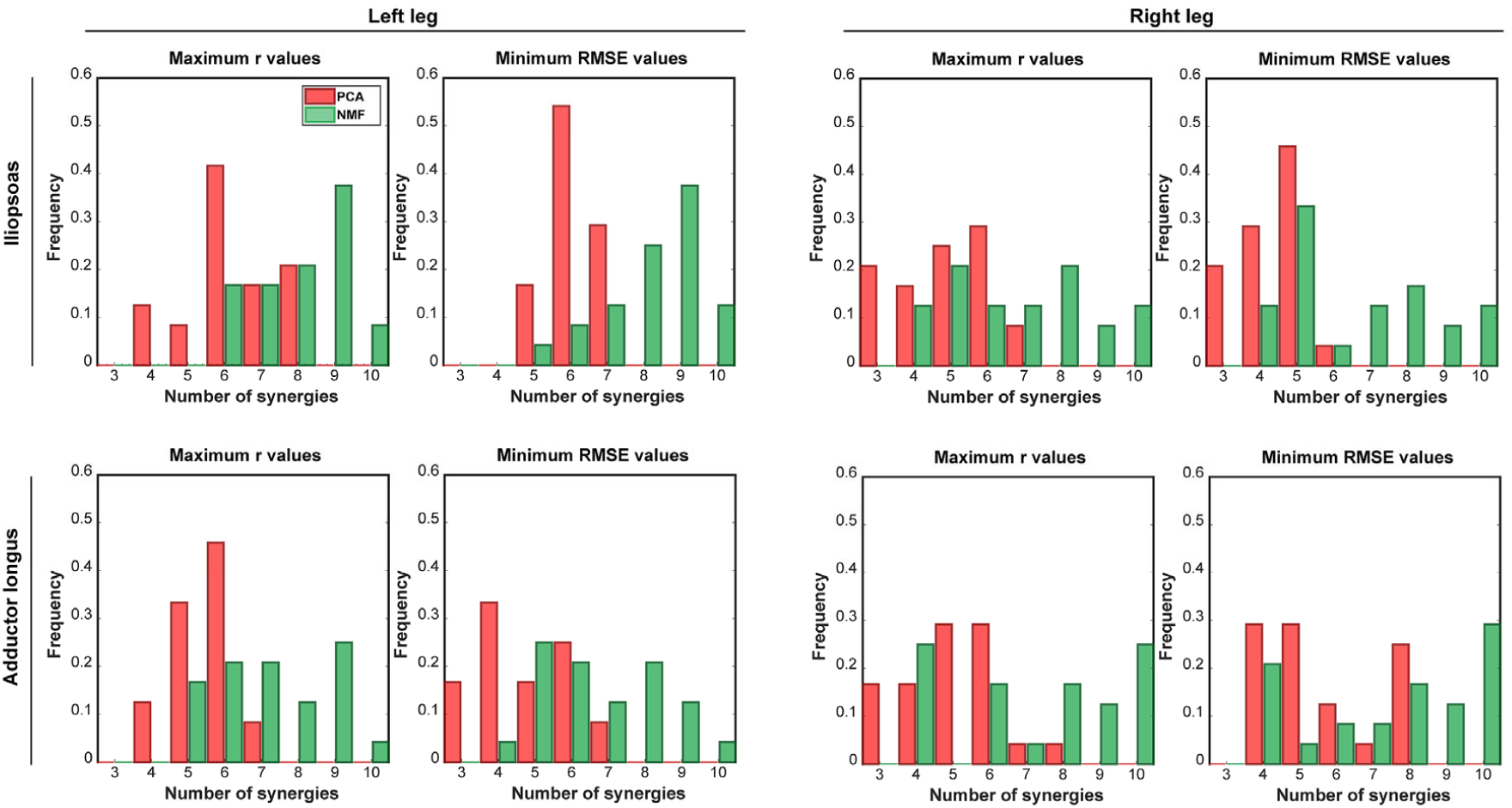
Distribution of the number of synergies that can provide maximum r values or minimum RMSE values across all calibration and evaluation trials with all the five EMG normalization methods. In each histogram, the horizontal axis reports the number of synergies, and the vertical axis shows the frequency with which the number of synergies can generate the best prediction of iliopsoas or adductor longus muscle excitations in terms of shape (indicated by r values) and magnitude (indicated by RMSE values). Red and green bars represent PCA-based and NMF-based synergy extrapolations, respectively. The right leg is paretic, and the left leg is non-paretic.

### 3.3 Joint moment prediction

Results of the three-factor ANOVA revealed that MAE values for joint moment matching were sensitive to both matrix factorization algorithm and number of synergies (*p* < 0.01) but insensitive to EMG normalization method (*p* = 0.12) for all muscle-leg-joint combinations. For the hipFE moment, MAE values for both PCA-based and NMF-based synergy extrapolations decreased as the number of synergies increased (*p* < 0.01). Nonetheless, PCA produced more accurate joint moment matching, as indicated by smaller MAE values, than did NMF (*p* < 0.01). In contrast, for the hipAA moment, neither number of synergies nor matrix factorization algorithm had a significant influence on MAE values. The one exception was adductor longus in the left leg, where MAE values decreased with an increasing number of synergies for both algorithms, but PCA had significantly smaller MAE values than NMF (*p* < 0.01) (Figs. 6 and 7).

**Fig. 6.**
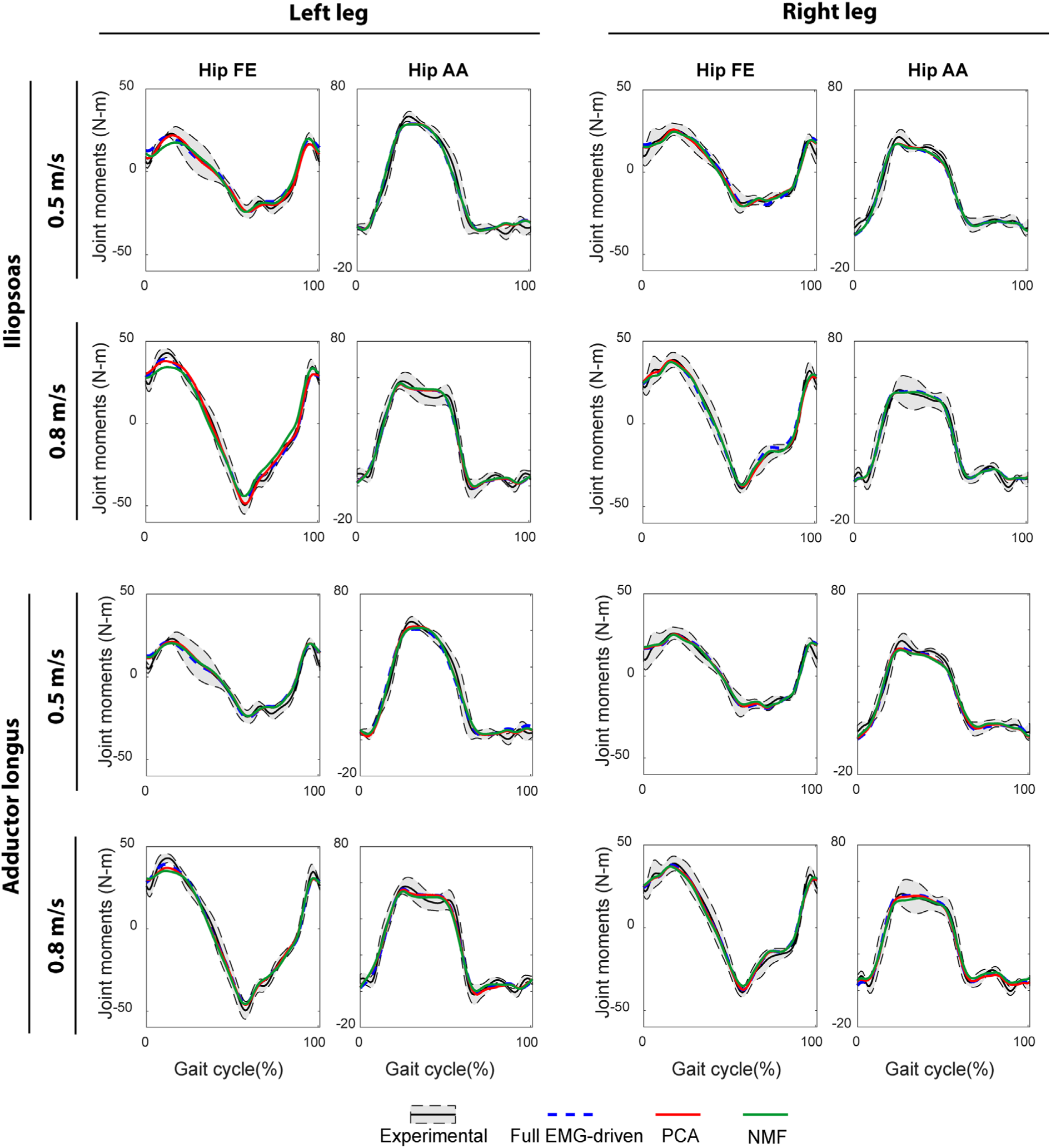
Average joint moment prediction across all calibration trials when performing full EMG-driven model calibration on step 1 (blue dash line) and synergy extrapolation on step 2 with PCA (red line) or NMF (green line), respectively. Black lines stand for the experimental joint moments from inverse dynamics, and the grey shaded areas represent ± 1 standard deviation. Six synergies are used for the generation of these representative results. Data is reported for the whole gait cycle with 0% being heel-strike and 100% being consecutive heel-strike events for both legs (right leg: paretic, left leg: non-paretic).

**Fig. 7.**
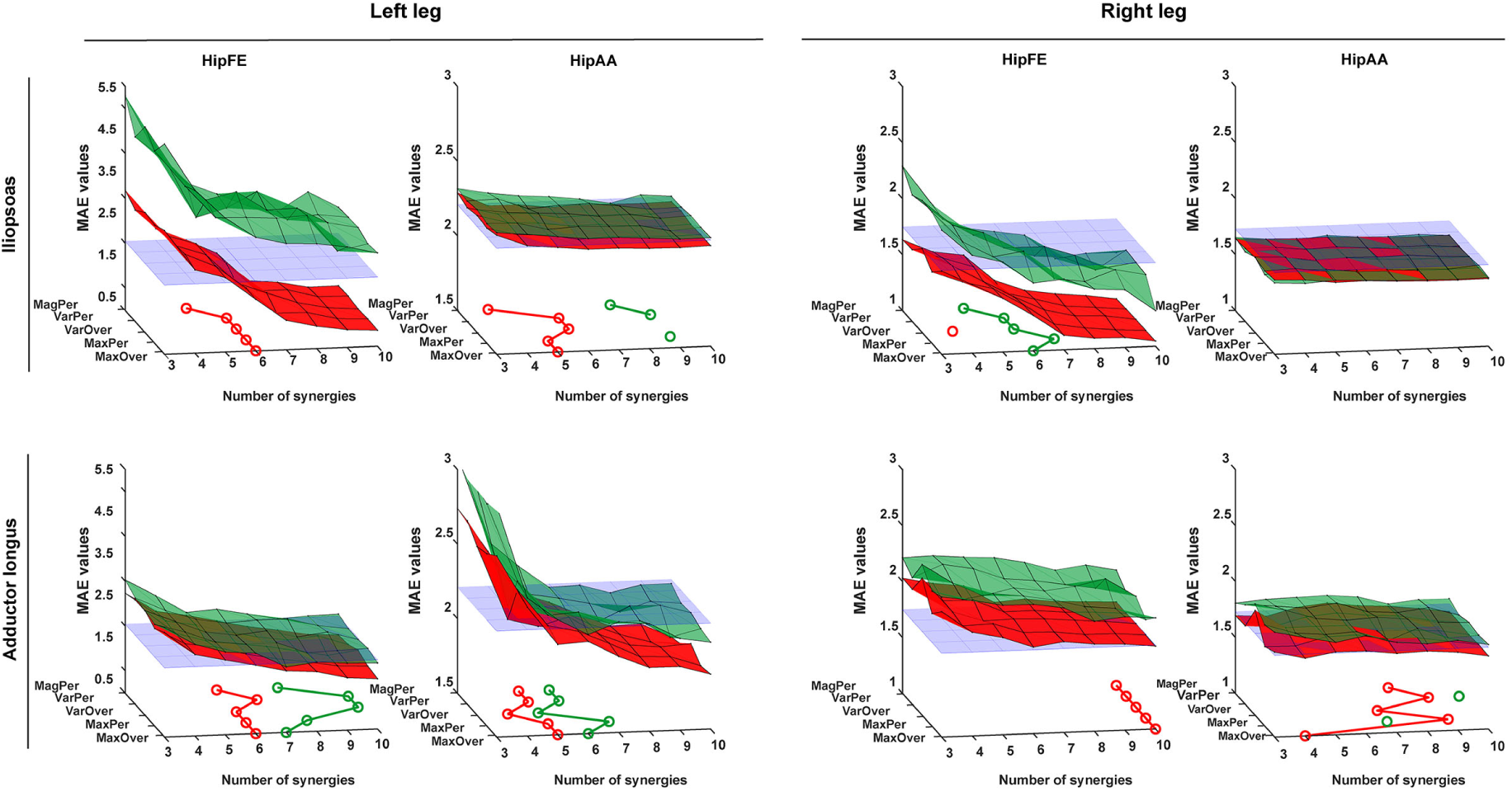
3D plots report the average MAE values for hip joint moment prediction across all trials by using 6 EMG normalization methods with an increase of the number of synergies from 3 to 10. The MAE values are averaged across all calibration and evaluation trials at 2 walking speeds for both legs (right leg: paretic, left leg: non-paretic). PCA- and NMF-based synergy extrapolations are indicated in red and green surfaces, respectively. The flat purple planes demonstrate the average MAE values for joint moment prediction with a full set of EMG signals at a specific leg. The circle lines on the bottom walls indicate where 3D surfaces of average MAE values for PCA- and NMF-based synergy extrapolation first intersect with the plane of average MAE for the full EMG-driven calibration, respectively, as the number of synergies increases. The absence of a circle means that there was no intersection corresponding to that synergy.

PCA-based and NMF-based synergy extrapolation led to different levels of joint moment tracking accuracy compared to the full EMG-driven calibration (Figs. 6 and 7). For example, for the hipFE moment with MaxOver, the surface comprising average MAE values for PCA-based synergy extrapolation intersected with the flat plane of average MAE for the full EMG-driven calibration at 6 synergies for iliopsoas in the left leg, 6 synergies for adductor longus in the left leg and 10 synergies for adductor longus in the right leg. In contrast, for the hipFE moment with MaxOver, the surfaces comprising MAE values for NMF-based synergy extrapolation only dropped below the plane of MAE values for the full EMG-driven calibration for iliopsoas in the right leg at 6 synergies and for adductor longus in the left leg at 7 synergies.

When *r* and RMSE values for muscle excitation matching were plotted as a function of MAE values for joint moment matching (Fig. 8), the observed trends were different for PCA-versus NMF-based synergy extrapolation. For PCA-based synergy extrapolation, the trends were parabolic, with a small region of MAE values corresponding to both largest *r* values and smallest RMSE values. In contrast, for NMF-based extrapolation, the observed trends were approximately linear, with the smallest MAE errors corresponding to the largest *r* values and smallest RMSE values. For both synergy extrapolation methods, MAE values for joint moment matching were approximately the same in the region where *r* values were the largest, and RMSE values were the smallest.

**Fig. 8.**
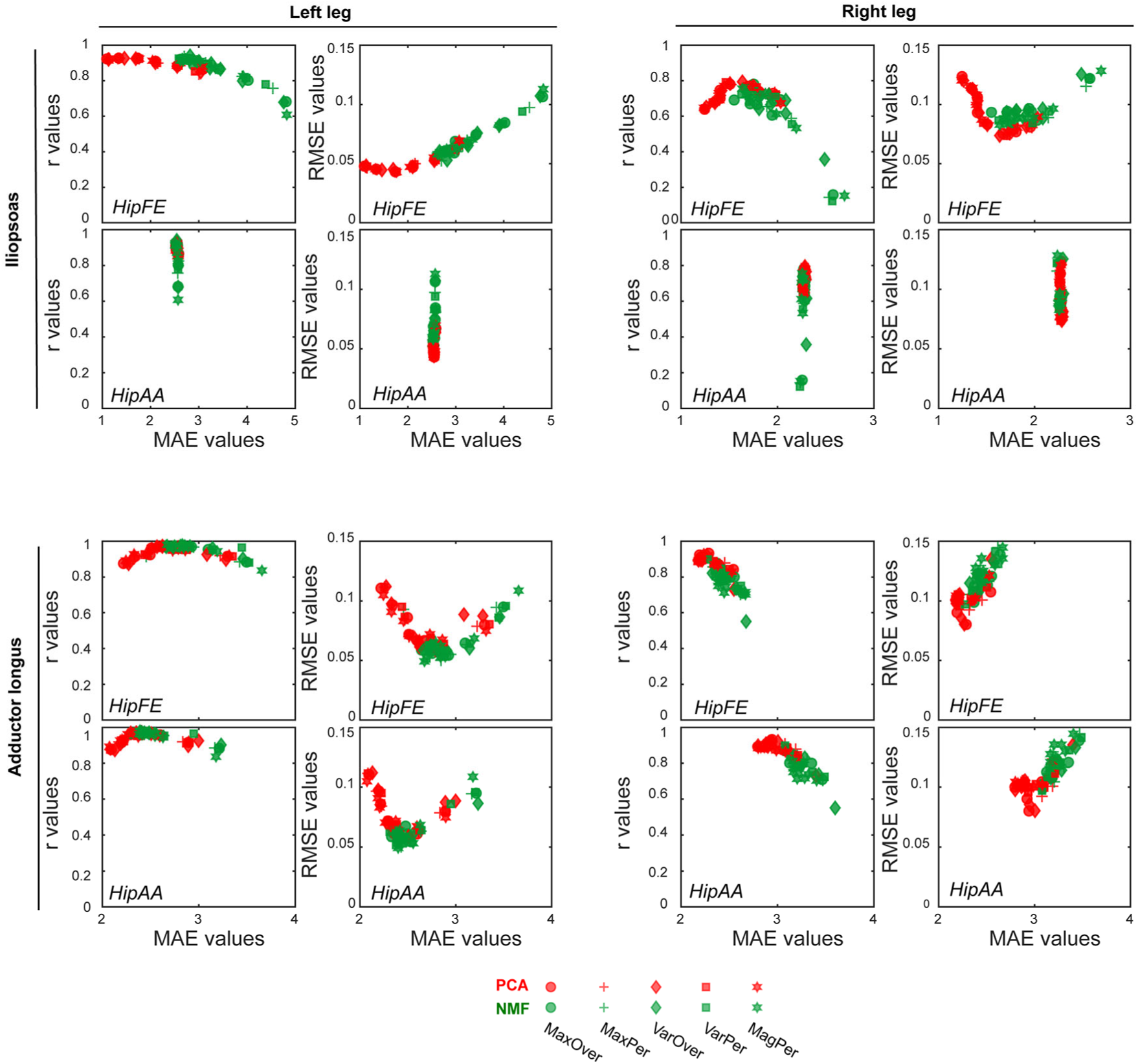
The trade-off relationship between the accuracy of joint moment tracking (indicated by MAE values in the horizontal axis) and accuracy of unmeasured muscle excitation reconstruction (indicated by r values or RMSE values in the y-axis) (red markers: PCA-based synergy extrapolation; green markers: NMF-based synergy extrapolation; 5 EMG normalization methods are represented by different marker shapes; the right leg is paretic, and the left leg is non-paretic).

## 4 Discussion

This study demonstrated that synergy extrapolation is able to predict unmeasured muscle excitations with reasonable reliability using a well-calibrated EMG-driven model. However, the reliability of the predictions was heavily influenced by the matrix factorization algorithm used and number of synergies, while EMG normalization method had little influence on synergy extrapolation results. Our results clearly showed that PCA was able to generate estimates of unmeasured muscle excitations that were more accurate in shape and magnitude, more robust to EMG normalization, and more consistent across all unmeasured muscle-leg combinations in comparison with NMF The results also highlighted that for PCA-based synergy extrapolation, a relatively low number of synergies, typically 5 or 6, always provided the most accurate predictions of unmeasured muscle excitations and joint moments simultaneously, while NMF was either not able to reproduce unmeasured muscle excitations with reasonable accuracy or required a large number of synergies, typically above 8, to attain a comparable outcome to PCA.

The synergy extrapolation method optimizes unmeasured synergy weights by tracking joint moments as closely as possible. We used the muscle synergy concept for the prediction of missing muscle excitations for several reasons. First, it is commonly theorized that muscle synergies are generated by the CNS to regulate highly redundant musculoskeletal systems in an efficient manner (Tresch et al., 2006; Torres-Oviedo and Ting, 2007). Second, the problem of finding unknown time-varying muscle excitations is reduced to the problem of finding a small number of synergy weights associated with unmeasured muscles, which significantly decreases the search space for the optimization. Third, there are no abrupt changes in predicted muscle excitations as is often observed with SO-based approaches, since predicted muscle excitations are linear combinations of weighted synergy excitations that are normally smooth. Fourth, by finding unknown synergy vector weights within the context of an EMG-driven model framework, the method is practically applicable in contrast to previous work that only demonstrated theoretical feasibility (Bianco et al., 2018). Sixth, synergy extrapolation results demonstrated that joint moments were matched as accurately as could be achieved by the full EMG-driven model (Fig. 6) (Meyer et al., 2017). Given these observations, the proposed synergy extrapolation method may outperform other EMG-driven muscle force estimators that either ignore deep muscles or use SO-based methods to estimate EMG signals for inaccessible muscles.

For the same number of synergies, PCA generally predicted unmeasured muscle excitations with noticeably greater accuracy than did NMF (Fig. 4). PCA and NMF are two of the most popular matrix factorization algorithms used for performing MSA. Although the MSA literature suggests that NMF generates synergy components that are highly correlated with those generated by PCA (Ivanenko et al., 2005; Cappellini et al., 2006; Banks et al., 2017), each of the algorithms decomposes a given dataset with different assumptions and constraints (Ting and Chvatal, 2010; Gallina et al., 2018). PCA is based on linear algebra that maintains orthogonality among all principal components during factorization, while NMF is a nonlinear optimization algorithm with potentially non-unique solutions in the non-negative space (Fig. 1 in the Supplementary Materials). PCA has been rejected by a few studies due to the inherent non-negative nature of muscle excitations (Ajiboye and Weir, 2009; Banks et al., 2017). However, our findings demonstrated several benefits of PCA over NMF for synergy extrapolation purposes. At least three observations help explain why PCA-based synergy extrapolation generally worked better than did NMF-based synergy extrapolation. First, for an equal number of synergies, PCA-derived components accounted for more variance in measured muscle excitations than did those derived using NMF, for 3 to 5 synergies in particular (Table 1). Our findings agree well with the results of variance accounted for by different numbers of synergies using when PCA and NMF are applied to high-density EMG data (Gallina et al., 2018). Second, PCA identifies synergy components that tend to describe the direction of the largest variance in the measured muscle excitations, with subsequent components being perpendicular to the previous ones, while NMF finds components that tend to represent the edge of a convex subspace in which all original EMG measurements lie (Ting and Chvatal, 2010; Lambert-Shirzad and Van der Loos, 2017). Furthermore, any of the data in the reduced-dimensionality space could be reconstructed due to negative and positive weights allowed by PCA. However, only data points representing unmeasured muscle excitations that were located within the defined subspace could be closely reproduced by NMF due to the constraints on non-negativity of the solutions (Tresch et al., 2006; Ting and Chvatal, 2010). Third, PCA-based synergy extrapolation had one more design variable - an offset - for each trial that represents the average value of the unmeasured muscle excitation, thereby adding one extra degree of freedom to the optimization problem. Therefore, better performance of PCA-based than NMF-based synergy extrapolation may be on account of the non-negativity constraints within NMF and the extra design variables for PCA, both of which make the feasible search space of NMF more restricted than that of PCA.

Synergy extrapolation performance was heavily influenced by the number of synergies used. Previous studies have reported that 3 to 6 synergies are adequate for reconstructing measured EMG signals with over 90% variance being accounted for during gait (Torres-Oviedo and Ting, 2007; Clark et al., 2009; Banks et al., 2017; Bianco et al., 2018) consistent with the observations in this study (Table 1). Not surprisingly, as the number of synergies increased, reconstruction accuracy of unmeasured muscle excitations using NMF increased before reaching a plateau, which was similar to reconstruction behavior for measured muscle excitations (Table 1). However, the change in PCA-based synergy extrapolation performance with an increasing number of synergies exhibited non-monotonic behavior (Fig. 4). Since synergy extrapolation iteratively predicted muscle excitations for iliopsoas or adductor longus through an optimization process, this additional flexibility likely allowed the optimizer to achieve lower joint moment tracking errors than those achieved by the full EMG-driven optimization. The same phenomenon can be observed for the hip flexion/extension moment using PCA (Fig.7). There, the synergy extrapolation performance peaked across different unmeasured muscle-leg combinations with the same number of synergies (Fig. 4). However, due to the non-negativity constraints built in NMF, the joint moment tracking errors stayed above or around the levels from the full EMG-driven optimization, which caused synergy extrapolation performance to level off when the number of synergies was above a certain number. Even though the variability existed in the number of synergies that generated the best synergy extrapolation results across trials, muscles, and legs, PCA needed fewer synergies than did NMF (Fig. 5), which simplifies the implementation of the synergy extrapolation method.

Though EMG normalization methods had no statistically significant influence on synergy extrapolation performance using PCA and NMF, the variability of average *r* and RMSE values across all five EMG normalization methods was considerably less for PCA than that for NMF using the same number of synergies (Fig. 4). PCA generates principal components that successively maximize variance (Tresch et al., 2006), which explains why each PCA-derived synergy excitation representing the corresponding variance for each primary component exhibited significant magnitude reduction (Supplementary Materials, Fig. 1). Additionally, the low-dimensional space in PCA defined by orthogonal principal components rotates as data are scaled differently across different muscles, but the rotated principal components are still highly descriptive of any points in the low-dimensional orthogonal space (Ting and Chvatal, 2010). However, NMF decomposition is based on assessing the quality of magnitude approximation, and scaling of original data differently across muscles would lead to deformation of the convex subspace that the components surround. With certain scaling schemes or normalization methods, the points accounting for unmeasured muscle excitations may not be able to stay in the NMF newly-generated subspace. Therefore, NMF-based synergy extrapolation performance would be sensitive to EMG normalization method, which is in agreement with the conclusions for NMF-based decomposition by Tresch et al., 2006. For different EMG normalization methods, the optimizer is presented with a more consistent searching space for PCA than for NMF, which could explain why PCA is less sensitive to EMG normalization method than is NMF. Therefore, PCA may eliminate the need to identify the true maximum muscle excitation for EMG normalization purposes in preparation for synergy extrapolation.

Another observation in this study was that, generally, for a given number of synergies, the non-paretic side (left leg) had higher reconstruction quality for both measured and unmeasured muscle excitations than the paretic side (right leg). For the same number of synergies, VAF values calculated for reconstructed measured muscle excitations in MSA were slightly higher for the non-paretic leg than the paretic leg (Table 1), which is not consistent with the findings reported by Clark et al., 2009. The underlying reason for this discrepancy could be the methodological differences between Clark et al.’s and our studies. For example, we performed MSA on a different set of muscles, which may have influenced the overall MSA outcomes (Banks et al., 2017) (Steele et al., 2013). Furthermore, the single subject post-stroke that participated in this study was high-functioning whereas Clark et al. studied 55 number of subjects; the heterogeneity in this clinical population may have also affected the conclusions. Our study intriguingly found that the reconstruction of unmeasured muscle excitations through synergy extrapolation was more accurate for the non-paretic side (left leg) than those for the paretic leg (right leg) (Fig.4), which is possibly due to less consistent EMG data of the healthy/non-paretic leg or more variable EMG signals across multiple trials for the paretic leg. For instance, the peaks appeared in the muscle excitations associated with adductor longus during the swing phase (60%-100% of gait cycle) were more drastic in the paretic side, and synergy extrapolation was unable to reproduce those as accurately it could do for the non-paretic side (Figs. 2 and 3). Additionally, to interpret the model calibration difference between legs, we observed that for the non-paretic side (left leg), as number of synergies increased, the joint moment tracking error for PCA-based synergy extrapolation started above and then dropped below the plane for the full EMG-driven calibration (Fig. 7). However, this behavior could not be seen in the paretic side (right leg), and the consequence of this phenomena was that none of the number of synergies could generate both muscle excitations and joint moments accurately.

Our study involved several important limitations that suggest areas for future investigation. First, we calibrated the EMG-driven model with a full set of EMG signals to obtain personalized model parameters (e.g., EMG scale factors, electromechanical delays, musculotendon parameter values, and geometric coefficients), and synergy extrapolation was feasible when the calibrated model parameters were held constant. In the future, a challenge will be to develop an optimization problem formulation where design variables defining both unmeasured synergy vector weights and EMG-driven model parameters are found simultaneously, the goal being to reproduce both joint moments and unmeasured muscle excitations accurately. Second, the extraction of measured muscle synergies was performed before the optimization process. However, since changing some parameter values (e.g., EMG scale factor) can change the synergy analysis outcome, MSA would need to be performed iteratively within optimization. Since PCA leads to unique and fast factorization solutions for synergy excitations while NMF does not (Tresch et al., 2006), PCA would be the preferred decomposition approach. This iterative decomposition might itself facilitate improvement in the estimation accuracy of unmeasured muscle excitations. Third, the synergy extrapolation framework was validated on cases where only one muscle (e.g. iliopsoas or adductor longus) was assumed to be missing. The approach should be evaluated further on cases where multiple muscles are assumed to be unmeasured concurrently. Further, this study used gait data from only one single subject post-stroke with extensive EMG data. Synergy extrapolation needs to be investigated in diverse subject populations, dynamic movement conditions, and experimental scenarios with other unmeasured muscles.

## 5 Conclusion

This study showed that synergy extrapolation is a viable option for estimating an unmeasured muscle excitation using synergy excitations extracted from measured muscle excitations. The study also demonstrated that methodological choices (i.e. matrix factorization algorithm and the number of synergies) made before MSA influence affect the accuracy with which unmeasured muscle excitations can be predicted. The synergy vector weights associated with unmeasured muscles were optimized by minimizing errors between the EMG-driven model predicted and experimental joint moments. Our results highlighted the fact that PCA was able to provide more accurate, reliable, and efficient estimates of unmeasured muscle excitations than was NMF. In general, PCA required 5 or 6 synergies to achieve the best prediction of unmeasured muscle excitations, and inclusion of additional synergies reduced synergy extrapolation performance. Better synergy extrapolation performance for PCA may be the result of the non-negativity restrictions imposed by NMF. Moreover, PCA was less sensitive to EMG normalization method than NMF, which may reduce the need to simulate the true maximum muscle excitations reliably for EMG normalization. Synergy extrapolation is of potential utility to address difficulties in collecting EMG signals from deep muscles inaccessible by surface electrodes, which is critical when predicting muscle forces with an EMG-driven musculoskeletal model. Our proposed synergy extrapolation method may facilitate the assessment of human neuromuscular control and biomechanics after rehabilitation or surgical treatment when EMG data collection is limited.

## 6 Acknowledgments

This work was conducted with support from the Cancer Prevention and Research Institute of Texas (CPRIT) RR170026.

## 7 Author Contributions Statement

CP and BF designed and performed the experiments; DA analyzed the data, prepared figures, and drafted the manuscript; MS and BF revised the manuscript; DA, MS, CP, and BF approved the final version of the manuscript.

## Reference

Ackermann, M., and van den Bogert, A. J. (2010). Optimality principles for model-based prediction of human gait. J. Biomech. 43, 1055–1060.

Ajiboye, A. B., and Weir, R. F. (2009). Muscle synergies as a predictive framework for the EMG patterns of new hand postures. J. Neural Eng. 6, 36004.

Allen, J. L., Kautz, S. A., and Neptune, R. R. (2013). The influence of merged muscle excitation modules on post-stroke hemiparetic walking performance. Clin. Biomech. 28, 697–704.

Allen, J. L., and Neptune, R. R. (2012). Three-dimensional modular control of human walking. J. Biomech. 45, 2157–2163.

Amarantini, D., and Martin, L. (2004). A method to combine numerical optimization and EMG data for the estimation of joint moments under dynamic conditions. J. Biomech. 37, 1393–1404. doi: https://doi.org/10.1016/j.jbiomech.2003.12.020.

Anderson, F. C., and Pandy, M. G. (2001). Static and dynamic optimization solutions for gait are practically equivalent. J. Biomech. 34, 153–161.

Arnold, E. M., Ward, S. R., Lieber, R. L., and Delp, S. L. (2010). A Model of the Lower Limb for Analysis of Human Movement. Ann. Biomed. Eng. 38, 269–279. doi: 10.1007/s10439-009-9852-5.

Banks, C. L., Pai, M. M., McGuirk, T. E., Fregly, B. J., and Patten, C. (2017). Methodological choices in muscle synergy analysis impact differentiation of physiological characteristics following stroke. Front. Comput. Neurosci. 11, 78.

Bianco, N. A., Patten, C., and Fregly, B. J. (2018). Can Measured Synergy Excitations Accurately Construct Unmeasured Muscle Excitations? J. Biomech. Eng. 140, 11011.

Bowden, M. G., Clark, D. J., and Kautz, S. A. (2010). Evaluation of abnormal synergy patterns poststroke: relationship of the Fugl-Meyer Assessment to hemiparetic locomotion. Neurorehabil. Neural Repair 24, 328–337.

Buchanan, T. S., Lloyd, D. G., Manal, K., and Besier, T. F. (2005). Estimation of muscle forces and joint moments using a forward-inverse dynamics model. Med. Sci. Sports Exerc. 37, 1911.

Cappellini, G., Ivanenko, Y. P., Poppele, R. E., and Lacquaniti, F. (2006). Motor patterns in human walking and running. J. Neurophysiol. 95, 3426–3437.

Clark, D. J., Ting, L. H., Zajac, F. E., Neptune, R. R., and Kautz, S. A. (2009). Merging of healthy motor modules predicts reduced locomotor performance and muscle coordination complexity post-stroke. J. Neurophysiol. 103, 844–857.

Contessa, P., and Luca, C. J. De (2013). Neural control of muscle force: indications from a simulation model. J. Neurophysiol. 109, 1548–1570.

Crowninshield, R. D., and Brand, R. A. (1981). A physiologically based criterion of muscle force prediction in locomotion. J. Biomech. 14, 793–801.

d’Avella, A., Saltiel, P., and Bizzi, E. (2003). Combinations of muscle synergies in the construction of a natural motor behavior. Nat. Neurosci. 6, 300–308.

Damsgaard, M., Rasmussen, J., Christensen, S. T., Surma, E., and De Zee, M. (2006). Analysis of musculoskeletal systems in the AnyBody Modeling System. Simul. Model. Pract. Theory 14, 1100–1111.

Del Vecchio, A., Ubeda, A., Sartori, M., Azorin, J. M., Felici, F., and Farina, D. (2018). Central nervous system modulates the neuromechanical delay in a broad range for the control of muscle force. J. Appl. Physiol. 125, 1404–1410.

Delp, S. L., Anderson, F. C., Arnold, A. S., Loan, P., Habib, A., John, C. T., et al. (2007). OpenSim: open-source software to create and analyze dynamic simulations of movement. IEEE Trans. Biomed. Eng. 54, 1940–1950.

Ebied, A., Kinney-Lang, E., Spyrou, L., and Escudero, J. (2018). Evaluation of matrix factorisation approaches for muscle synergy extraction. Med. Eng. Phys. 57, 51–60. doi: https://doi.org/10.1016/j.medengphy.2018.04.003.

Falisse, A., Pitto, L., Kainz, H., Hoang, H., Wesseling, M., Van Rossom, S., et al. (2020). Physics-based simulations to predict the differential effects of motor control and musculoskeletal deficits on gait dysfunction in cerebral palsy: a retrospective case study. Front. Hum. Neurosci. 14, 40.

Farina, D., Cescon, C., and Merletti, R. (2002). Influence of anatomical, physical, and detection-system parameters on surface EMG. Biol. Cybern. 86, 445–456.

Fregly, B. J., Besier, T. F., Lloyd, D. G., Delp, S. L., Banks, S. A., Pandy, M. G., et al. (2012a). Grand challenge competition to predict in vivo knee loads. J. Orthop. Res. 30, 503–513.

Fregly, B. J., Boninger, M. L., and Reinkensmeyer, D. J. (2012b). Personalized neuromusculoskeletal modeling to improve treatment of mobility impairments: a perspective from European research sites. J. Neuroeng. Rehabil. 9, 18.

Gallina, A., Garland, S. J., and Wakeling, J. M. (2018). Identification of regional activation by factorization of high-density surface EMG signals: A comparison of Principal Component Analysis and Non-negative Matrix factorization. J. Electromyogr. Kinesiol. 41, 116–123. doi: https://doi.org/10.1016/j.jelekin.2018.05.002.

He, J., Levine, W. S., and Loeb, G. E. (1991). Feedback gains for correcting small perturbations to standing posture. IEEE Trans. Automat. Contr. 36, 322–332. doi: 10.1109/9.73565.

Heintz, S., and Gutierrez-Farewik, E. M. (2007). Static optimization of muscle forces during gait in comparison to EMG-to-force processing approach. Gait Posture 26, 279–288.

Hill, A. V. (1938). The heat of shortening and the dynamic constants of muscle. Proc. R. Soc. London. Ser. B-Biological Sci. 126, 136–195.

Hug, F., Turpin, N. A., Dorel, S., and Guével, A. (2012). Smoothing of electromyographic signals can influence the number of extracted muscle synergies. Clin. Neurophysiol. Off. J. Int. Fed. Clin. Neurophysiol. 123, 1895.

Ivanenko, Y. P., Cappellini, G., Dominici, N., Poppele, R. E., and Lacquaniti, F. (2005). Coordination of locomotion with voluntary movements in humans. J. Neurosci. 25, 7238–7253.

Kristiansen, M., Madeleine, P., Hansen, E. A., and Samani, A. (2015). Inter-subject variability of muscle synergies during bench press in power lifters and untrained individuals. Scand. J. Med. Sci. Sports 25, 89–97.

Kumar, D., Rudolph, K. S., and Manal, K. T. (2012). EMG-driven modeling approach to muscle force and joint load estimations: Case study in knee osteoarthritis. J. Orthop. Res. 30, 377–383.

Lambert-Shirzad, N., and Van der Loos, H. F. M. (2017). On identifying kinematic and muscle synergies: a comparison of matrix factorization methods using experimental data from the healthy population. J. Neurophysiol. 117, 290–302.

Lee, D. D., and Seung, H. S. (1999). Learning the parts of objects by non-negative matrix factorization. Nature 401, 788–791.

Lloyd, D. G., and Besier, T. F. (2003). An EMG-driven musculoskeletal model to estimate muscle forces and knee joint moments in vivo. J. Biomech. 36, 765–776. doi: https://doi.org/10.1016/S0021-9290(03)00010-1.

Manal, K., and Buchanan, T. S. (2003). A one-parameter neural activation to muscle activation model: estimating isometric joint moments from electromyograms. J. Biomech. 36, 1197–1202. doi: https://doi.org/10.1016/S0021-9290(03)00152-0.

McLean, S. G., Huang, X., and van den Bogert, A. J. (2005). Association between lower extremity posture at contact and peak knee valgus moment during sidestepping: implications for ACL injury. Clin. Biomech. 20, 863–870.

Mehrabi, N., Schwartz, M. H., and Steele, K. M. (2019). Can altered muscle synergies control unimpaired gait? J. Biomech. 90, 84–91.

Mehryar, P., Shourijeh, M. S., Rezaeian, T., Khandan, A. R., Messenger, N., O’Connor, R., et al. (2020). Differences in muscle synergies between healthy subjects and transfemoral amputees during normal transient-state walking speed. Gait Posture 76, 98–103, doi: https://doi.org/10.1016/j.gaitpost.2019.10.034

Menegaldo, L. L., de Toledo Fleury, A., and Weber, H. I. (2004). Moment arms and musculotendon lengths estimation for a three-dimensional lower-limb model. J. Biomech. 37, 1447–1453.

Meyer, A. J., Eskinazi, I., Jackson, J. N., Rao, A. V, Patten, C., and Fregly, B. J. (2016a). Muscle synergies facilitate computational prediction of subject-specific walking motions. Front. Bioeng. Biotechnol. 4, 1–26.

Meyer, A. J., Eskinazi, I., Jackson, J. N., Rao, A. V, Patten, C., and Fregly, B. J. (2016b). Muscle synergies facilitate computational prediction of subject-specific walking motions. Front. Bioeng. Biotechnol. 4, 77.

Meyer, A. J., Patten, C., and Fregly, B. J. (2017). Lower extremity EMG-driven modeling of walking with automated adjustment of musculoskeletal geometry. PLoS One 12, e0179698.

Oliveira, A. S., Gizzi, L., Farina, D., and Kersting, U. G. (2014). Motor modules of human locomotion: influence of EMG averaging, concatenation, and number of step cycles. Front. Hum. Neurosci. 8, 335.

Olree, K. S., and Vaughan, C. L. (1995). Fundamental patterns of bilateral muscle activity in human locomotion. Biol. Cybern. 73, 409–414.

Péter, A., Andersson, E., Hegyi, A., Finni, T., Tarassova, O., Cronin, N., et al. (2019). Comparing surface and fine-wire electromyography activity of lower leg muscles at different walking speeds. Front. Physiol. 10, 1283.

Pitto, L., Kainz, H., Falisse, A., Wesseling, M., Van Rossom, S., Hoang, H., et al. (2019). SimCP: A Simulation Platform to Predict Gait Performance Following Orthopedic Intervention in Children With Cerebral Palsy. Front. Neurorobotics 13, 54. Available at: https://www.frontiersin.org/article/10.3389/fnbot.2019.00054.

Racinais, S., Maffiuletti, N. A., and Girard, O. (2013). M-wave, H-and V-reflex recruitment curves during maximal voluntary contraction. J. Clin. Neurophysiol. 30, 415–421.

Reinbolt, J. A., Schutte, J. F., Fregly, B. J., Koh, B. Il, Haftka, R. T., George, A. D., et al. (2005). Determination of patient-specific multi-joint kinematic models through two-level optimization. J. Biomech. 38, 621–626. doi: https://doi.org/10.1016/j.jbiomech.2004.03.031.

Ruiz Garate, V., Parri, A., Yan, T., Munih, M., Molino Lova, R., Vitiello, N., et al. (2017). Experimental Validation of Motor Primitive-Based Control for Leg Exoskeletons during Continuous Multi-Locomotion Tasks. Front. Neurorobotics 11, 15. Available at: https://www.frontiersin.org/article/10.3389/fnbot.2017.00015.

Shourijeh, M., and McPhee, J. (2014). Forward Dynamic Optimization of Human Gait Simulations: A Global Parameterization Approach. ASME J. Comput. Nonlinear Dyn. 9, 31018. doi: https://doi.org/10.1115/1.4026266.

Sartori, M., Farina, D., and Lloyd, D. G. (2014). Hybrid neuromusculoskeletal modeling to best track joint moments using a balance between muscle excitations derived from electromyograms and optimization. J. Biomech. 47, 3613–3621.

Sartori, M., Reggiani, M., Farina, D., and Lloyd, D. G. (2012). EMG-Driven Forward-Dynamic Estimation of Muscle Force and Joint Moment about Multiple Degrees of Freedom in the Human Lower Extremity. PLoS One 7, e52618. Available at: https://doi.org/10.1371/journal.pone.0052618.

Sauder, N. R., Meyer, A. J., Allen, J. L., Ting, L. H., Kesar, T. M., and Fregly, B. J. (2019). Computational Design of FastFES Treatment to Improve Propulsive Force Symmetry During Post-stroke Gait: A Feasibility Study. Front. Neurorobotics 13, 80. Available at: https://www.frontiersin.org/article/10.3389/fnbot.2019.00080.

Seth, A., Hicks, J. L., Uchida, T. K., Habib, A., Dembia, C. L., Dunne, J. J., et al. (2018). OpenSim: Simulating musculoskeletal dynamics and neuromuscular control to study human and animal movement. PLoS Comput. Biol. 14.

Shao, Q., Bassett, D. N., Manal, K., and Buchanan, T. S. (2009). An EMG-driven model to estimate muscle forces and joint moments in stroke patients. Comput. Biol. Med. 39, 1083–1088. doi: https://doi.org/10.1016/j.compbiomed.2009.09.002.

Shourijeh, M. S., Flaxman, T. E., and Benoit, D. L. (2016). An approach for improving repeatability and reliability of non-negative matrix factorization for muscle synergy analysis. J. Electromyogr. Kinesiol. 26, 36–43, doi: https://doi.org/10.1016/j.jelekin.2015.12.001

Shourijeh, M. S., and Fregly, B. J. (2020). Muscle Synergies Modify Optimization Estimates of Joint Stiffness During Walking. J. Biomech. Eng. 142, 011011. doi: https://doi.org/10.1115/1.4044310

Shuman, B. R., Schwartz, M. H., and Steele, K. M. (2017). Electromyography Data Processing Impacts Muscle Synergies during Gait for Unimpaired Children and Children with Cerebral Palsy. Front. Comput. Neurosci. 11, 50. Available at: https://www.frontiersin.org/article/10.3389/fncom.2017.00050.

Steele, K. M., Tresch, M. C., and Perreault, E. J. (2013). The number and choice of muscles impact the results of muscle synergy analyses. Front. Comput. Neurosci. 7, 105.

Taylor, R. (1990). Interpretation of the correlation coefficient: a basic review. J. diagnostic Med. Sonogr. 6, 35–39.

Thelen, D. G., Anderson, F. C., and Delp, S. L. (2003). Generating dynamic simulations of movement using computed muscle control. J. Biomech. 36, 321–328.

Ting, L. H., and Chvatal, S. A. (2010). Decomposing muscle activity in motor tasks. Mot. Control Theor. Exp. Appl. Oxf. Univ. Press. New York, 102v – 138.

Ting, L. H., and Macpherson, J. M. (2005). A limited set of muscle synergies for force control during a postural task. J. Neurophysiol. 93, 609–613.

Torres-Oviedo, G., and Ting, L. H. (2007). Muscle synergies characterizing human postural responses. J. Neurophysiol. 98, 2144–2156.

Tresch, M. C., Cheung, V. C. K., and d’Avella, A. (2006). Matrix factorization algorithms for the identification of muscle synergies: evaluation on simulated and experimental data sets. J. Neurophysiol. 95, 2199–2212.

Tresch, M. C., Saltiel, P., and Bizzi, E. (1999). The construction of movement by the spinal cord. Nat. Neurosci. 2, 162–167.

Walter, J. P., Kinney, A. L., Banks, S. A., D’Lima, D. D., Besier, T. F., Lloyd, D. G., et al. (2014). Muscle synergies may improve optimization prediction of knee contact forces during walking. J. Biomech. Eng. 136, 21031.

Zajac, F. E. (1989). Muscle and tendon: properties, models, scaling, and application to biomechanics and motor control. Crit. Rev. Biomed. Eng. 17, 359–411.

Zonnino, A., and Sergi, F. (2019). Model-based estimation of individual muscle force based on measurements of muscle activity in forearm muscles during isometric tasks. IEEE Trans. Biomed. Eng., 1. doi: 10.1109/TBME.2019.2909171.

