## Supplemental Material for "Evaluation of Synergy Extrapolation for Predicting Unmeasured Muscle Excitations from Measured Muscle Synergies" (from another version of this preprint: version 1, 2020-08-06)

### *Supplementary Material*

#### 1 Muscles in each leg

**Table 1** List of the muscles in the EMG-driven model and corresponding degrees of freedom (DOFs).

| EMG channel | EMG label | EMG type | DOFs | Muscle |
|---|---|---|---|---|
| Adductor longus | AddLong | Fine wire | Hip FE/Hip AA | Adductor brevis |
| | | | | Adductor longus |
| | | | | Adductor magnus distal |
| | | | | Adductor magnus ischial |
| | | | | Adductor magnus middle |
| | | | | Adductor magnus proximal |
| Gluteus maximus | GlutMax | Surface | | Gluteus maximus superior |
| | | | | Gluteus maximus middle |
| | | | | Gluteus maximus inferior |
| Gluteus medius | GlutMed | Surface | | Gluteus medius anterior |
| | | | | Gluteus medius middle |
| | | | | Gluteus medius posterior |
| | | | | Gluteus minimus anterior |
| | | | | Gluteus minimus middle |
| | | | | Gluteus minimus posterior |
| Iliacus or Psoas | Iliopsoas | Fine wire | | Iliacus |
| | | | | Psoas |
| Semimembranosus | SemiMemb | Surface | Hip FE/Hip AA/Knee FE | Semimembranosus |
| | | | | Semitendinosus |
| Biceps femoris long head | BicFemLong | Surface | Hip FE/Hip AA/Knee FE | Biceps femoris long head |
| | | | Knee FE | Biceps femoris short head |
| Rectus femoris | RecFem | Surface | Hip FE/Hip AA/Knee FE | Rectus femoris |
| Vastus medialis | VastMed | Surface | Knee FE | Vastus medialis |
| | | | | Vastus intermedius |
| Vastus lateralis | VastLat | Surface | | Vastus lateralis |
| Medial gastrocnemius | GasMed | Surface | Knee FE/Ankle PDF/Ankle IE | Lateral gastrocnemius |
| | | | | Medial gastrocnemius |
| Tibialis anterior | TibAnt | Surface | Ankle PDF/Ankle IE | Tibialis anterior |
| Tibialis posterior | TibPost | Fine wire | | Tibialis posterior |
| Peroneus longus | PeroneusLong | Surface | | Peroneus brevis |
| | | | | Peroneus longus |
| | | | | Peroneus tertius |
| Soleus | Sol | Surface | | Soleus |
| Extensor digitorum longus | ExtDigLong | Fine wire | | Extensor digitorum longus |
| Flexor digitorum Longus | FlexDigLong | Fine wire | | Flexor digitorum Longus |

Hip FE: hip flexion/extension; Hip AA: hip adduction/abduction; Knee FE: knee flexion/extension; Ankle PDF: ankle plantarflexion/dorsiflexion; Ankle IE: ankle inversion/eversion.

#### 2 Muscle synergy analysis

Representative synergy components identified by PCA (Figure 1 (A)) composed of synergy excitations and corresponding weights that were both positive and negative. The percentage of variance that each synergy component explained decreased from 64% to 1% from the first to the sixth synergy, where the magnitude of each corresponding synergy excitation also experienced a significant drop. In contrast to PCA, the synergy excitations and weights determined by NMF (Figure 1 (B)) contained only non-negative values as constrained by the algorithm itself, and the variance that each component accounted for stayed at comparable levels. All the NMF-identified synergy excitations also had comparable magnitudes.

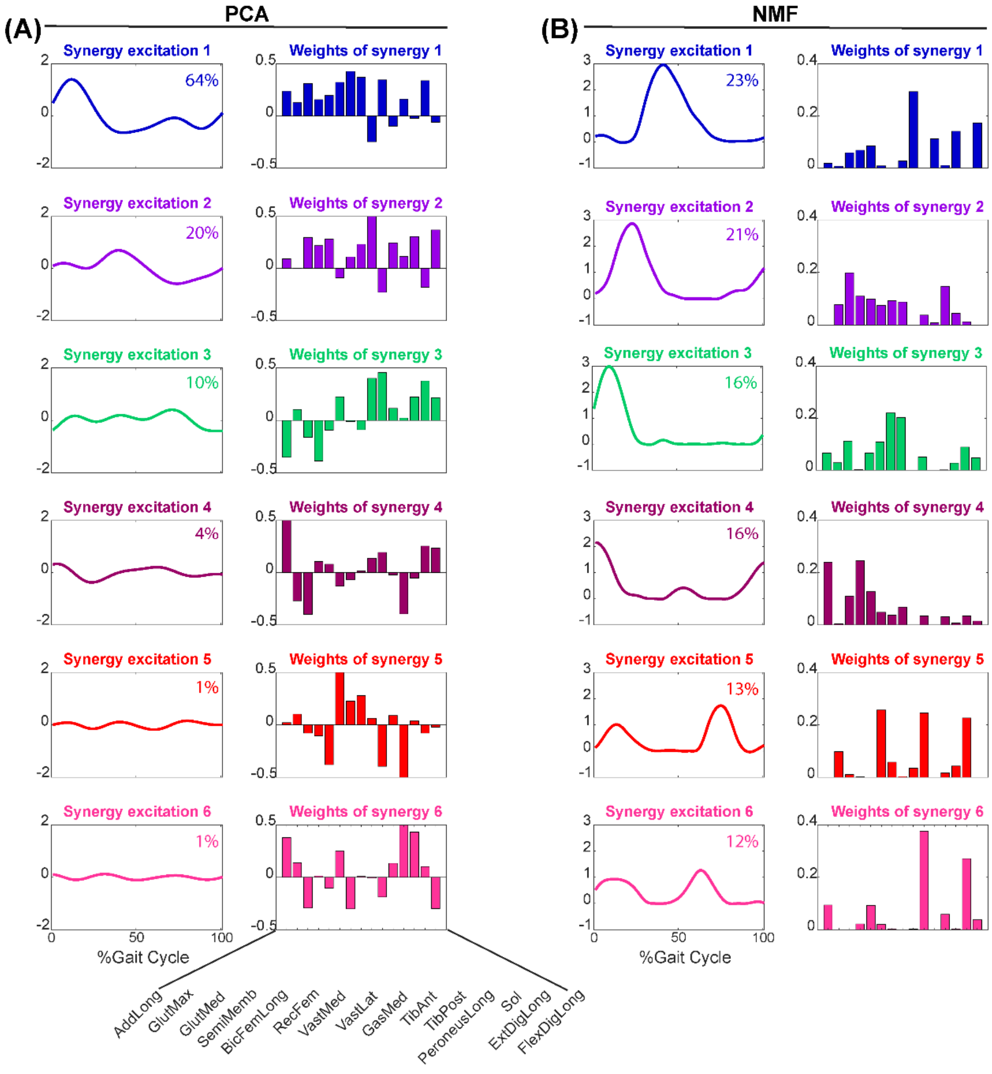


**Figure 1** Representative results of synergy excitations and corresponding synergy weights from measured EMGs identified using (A) PCA and (B) NMF, respectively. The sample data set was collected from the non-paretic side (left) of the patient when walking at his self-selected speed (0.5 m/s). 'Iliopsoas' EMG channel was assumed to be unmeasured, and the remaining 15 EMG channels were normalized to maximum values over all trials (MaxOver) before being decomposed for this case. Percentages represent the amount of total data VAF contributed by each synergy excitation, and the synergy components were sorted according to the descending percentage of VAF.
